# Pericyte and Endothelial Primary Cilia and Centrioles have Disparate Organization Across the Brain Microvasculature

**DOI:** 10.1101/2025.11.19.689283

**Authors:** Søren Grubb, Vanshika Chaddha, Jennifer Lippincott-Schwartz, Carolyn M. Ott, Amreen Mughal

**Affiliations:** Center for Translational Neuromedicine, Faculty of Health Sciences, University of Copenhagen, DK-2200 Copenhagen N, Denmark; The Neurovascular Research Unit, Stroke Branch-National Institute of Neurological Disorders and Stroke, Translational Vascular Medicine Branch-National Heart, Lung, and Blood Institute, National Institutes of Health, Bethesda, MD, USA; Janelia Research Campus, Howard Hughes Medical Institute, Ashburn, VA 20147, USA

**Keywords:** Brain Vasculature, Primary Cilia, Centrioles, Blood Flow, Neurovascular Coupling, Astrocytes, Neurons, Cell Polarity, Endothelial Cells, Mural Cells, Pericytes, Capillaries, Arterioles, Venules

## Abstract

The brain microvascular functions are strongly influenced by the local microenvironment and cellular organization. Intracellular organelles, including primary cilia and centrioles, play critical roles in sensing and transmitting environmental cues and maintaining vascular integrity. However, their distribution across the brain vasculature remains poorly understood. In this study, we utilized publicly available large-volume electron microscopy datasets encompassing the cerebral vasculature from pial arterioles through parenchymal capillaries to pial venules. We systematically analyzed the cellular organization and characterized the distribution of primary cilia and centrioles in the mouse and human brain microvasculature. We found primary cilia exclusively on human cortical endothelial cells (ECs), indicating inter-species differences between mouse and human. Primary cilia were frequently present on mural cells (MCs, smooth muscle cells or pericytes) surrounding venules and capillaries but rarely observed on arterioles in both mouse and human brains. These MC primary cilia exhibited heterogeneity in ciliogenesis, including cells with ciliary pockets, surface cilia, and a hybrid configuration we refer as a partial pocket. In the mouse brain, many MC primary cilia were closely ensheathed by astrocytic endfeet and occasionally extended between them to establish proximity to synapses, whereas all primary cilia in the human brain were confined within the basal lamina. Our analysis of cellular density revealed similar EC densities between arterioles and venules in mice, but not in human. EC centrioles were consistently positioned against the direction of blood flow relative to the nuclei, suggesting that they may serve as a structural marker for flow direction. Collectively, these findings provide a comprehensive characterization of primary cilia and centrioles, highlighting distinct interspecies differences between mouse and human brain microvasculature. The proximity to neural cells and gradient distribution of these subcellular structures suggest that they may act as antennae for sensing mechanical and chemical signals within the brain microvascular environment.

## Introduction

The brain is a computationally intensive and metabolically demanding organ, where 20% of body energy gets consumed. The tight regulation of nutrient supply to the active brain region is achieved through close communication between neural cells and blood vessels, known as neurovascular coupling (NVC). For NVC to work efficiently, the brain has a tightly woven network of blood vessels with varying caliber, cellular organization and molecular machinery. Although participation of different vascular cells has been described in NVC and is the basis for several brain imaging modalities, we are still lacking a thorough understanding of cellular and subcellular organization of the microvasculature.

Pial arterioles carry blood across the brain surface within the pia mater before descending into the brain parenchyma as penetrating arterioles, which further branch into an extensive capillary network. These capillaries diverge from arterioles into 3-6 branches, then re-converge through 3-6 branches before draining into ascending venules that return blood to the surface pial venules ^1–3^. The microvascular network is in close proximity to neural cells, and vessel diameters range from approximately 3-8 µm at the capillary level, 10-50 µm for penetrating arterioles and ascending venules, and 50-100 µm for larger pial arterioles and venules ^4,5^.

The cellular and subcellular organization of blood vessels is critical for regulating cerebral blood flow (CBF) ^1,5^. All blood vessels possess an innermost layer of endothelial cells (ECs), followed by mural cells (MCs). These ECs and MCs work in the close coordination to regulate the vascular tone ^6^ and have functional implications in control of cerebral hemodynamics, autoregulation, and NVC, to list a few. MCs comprise both smooth muscle cells (SMCs) and pericytes, and their distribution varies across the vasculature (detailed review ^1^). Single or multiple layers of SMCs encircle arterioles, while capillaries have gradient distribution of pericytes and venules are thought to have limited or no MC coverage ^7–9^. Given the recent findings about the functional role of the venous vasculature in the NVC ^10^ and aging associated pathophysiological changes in the venous microvasculature ^11^, understanding the position and organization of cellular and subcellular structures specifically in the venous vasculature is foundational to understanding their role in NVC among other functions.

Within ECs and MCs, centrosomes are key subcellular structures that create and contribute to cell polarization and potentially to endothelial cell barrier function ^12,13^. Each centrosome generally includes two centrioles, cylindrical array of microtubule triplets, one of which, the mother centriole, has specialized appendages that allow it to serve as the origin for a primary cilium. Cilia are the sensory organelles populated by specifically trafficked receptors able to detect changes in extracellular signaling molecules including proteins, lipids and ions ^14^. Cilia formation is thought to originate either from a centriole docked at the plasma membrane, generating a surface cilium, or by recruitment of vesicles to the mother centriole distal appendages ^15^. In the latter case the nascent cilium grows enveloped by a vesicle that eventually fuses with the plasma membrane and the membrane that had been enveloping the cilium is converted into a ciliary pocket ^16,17^. Surface cilia differ from pocket cilia in that their entire ciliary membrane is outside the cell-more fully exposed to the extracellular environment.

The functions of cilia in vascular cells have not been fully established. Mutations affecting the cilia localized receptor, Notch3 ^18^, cause Cerebral Autosomal Dominant Arteriopathy with Subcortical Infarcts and Leukoencephalopathy (CADASIL), a genetic form of vascular cognitive impairment and dementia (VCID) ^19,20^. Across the vasculature, cilia have been reported in mouse, rat, bovine, rabbit, zebrafish human brain and peripheral ECs ^21–32^. However, most of these studies are performed in early development stages and in cultured cells, making it difficult to determine if cilia are present in vascular endothelial cells in adults. A recent study showed that the majority of cilia are lost with endothelial differentiation during embryogenesis and only 4% lymphatic ECs have primary cilia in the adult mice ^33^. In line with ECs, our knowledge about the presence of cilia in brain mural cells (smooth muscle cells/pericytes) across the entire length of microvasculature remains limited ^34–39^.

Traditional approaches to study cilia and centrosome organization, such as confocal microscopy of immunostained brain sections, do not capture the full spatial context of the microvascular network. Recent advances in large-volume electron microscopy (EM) datasets with high subcellular resolution have made it possible to locate cilia and centrosomes across hundreds of neural cells ^40–42^. Moreover, these datasets allowed identification of microglial contact to the vasculature ^43^, ultrastructure of precapillary sphincters ^44^ and peg-socket interactions between pericyte-endothelial cell ^45,46^. To map primary cilia and centrosomes across the microvasculature, we first identified ECs and MCs in the vasculature from pial and penetrating arterioles through capillaries to pial and ascending venules. We then located cilia and centrosomes and assessed their locations, orientations, and cellular or extracellular environment. Our analyses both uncover differences in ciliary and centrosomal architecture between species and across the vascular network and provide a foundation for future mechanistic studies on their role in NVC and other brain vascular functions.

## Methods

### Large volume EM datasets

We utilized three publicly available large volume EM datasets: mouse visual cortex (MICrONS ^47^), mouse hippocampus (CA1 region ^48^), and human cerebral cortex (h01 ^49^ along with some small datasets (**Table 1**). The MICrONS Consortium datasets, mouse heart, choroid plexus and visual cortex and h01 datasets were visualized and quantified using Neuroglancer, an open-source browser-based viewer (https://www.microns-explorer.org/visualization), whereas the CA1, Macaque L2/3 S1 and STG dataset were visualized and quantified with Webknossos, an open-source tool (https://home.webknossos.org/).

**Table 1.** Attributes of public 3D-EM datasets from brain tissue.

| Source | Species | Brain Region(s) | Volume |
| --- | --- | --- | --- |
| MICrONS Consortium<br>Cubic MM<sup>3</sup> <sup>47</sup> | Mouse (P87) | Primary visual cortex and higher visual areas across all cortical layers | Volume: 1 mm<sup>3</sup> (1.3 x 0.87 x 0.82 mm)<br>Resolution: 4 x 4 x 40 nm per voxel |
| hippocampal/CA1 dataset <sup>48</sup> | Mouse (P29 female) | All layers of the CA1 region | Volume: 0.28 mm<sup>3</sup> (1.1 x 0.92 x 0.143 mm)<br>Resolution: 4 x 4 x 35 nm per voxel |
| h01 dataset <sup>49</sup> | Human (45-year-old woman) | Cerebral cortex, temporal lobe | Volume: 1 mm<sup>3</sup> (3 x 2 x 0.15 mm)<br>Resolution: 4 x 4 x 33.9 nm per voxel |
| MICrONS L2/3<br>https://layer23.microns- | Mouse (P36) | Primary visual cortex (Layer 1, 2/3) | Volume: 250 x 140 x 90 μm<br>Resolution: 3.58 x 3.58 x 40 nm per |
| explorer.org/ |  |  | voxel |
| Lee et al.<sup>50</sup> | Mouse (C57BL/6, 9 months, Male) | Primary visual cortex (Layer 1, 2/3) | Volume: 450 x 450 x 150 $\mu\text{m}$<br>Resolution: 4 x 4 x 40 nm per voxel |
| Macaque L2/3 S1 and STG<sup>51</sup> | Macaque (15.6 old, right-handed, male) | Primary somatosensory cortex (Superior temporal gyrus (STG) and S1 region) | Volume: 176x 224 x 100 $\mu\text{m}$ (S1)<br>175 x 225 x 108 $\mu\text{m}$ (STG)<br>Resolution: 11.24 x 11.24 x 28-30 nM voxel size |
| Mouse Heart<br>(https://doi.org/10.25378/janelia.24020919.v1) | Mouse (P7), C57BL/6J from Jackson Lab | Heart tissue | Volume: 153 x 151 x 136 $\mu\text{m}$<br>Resolution: 8 x 8 x 8 nM voxel size |
| Choroid plexus<br>(https://doi.org/10.1101/2025.01.21.633415) | Mouse (P14 brain), C57BL/6J from Jackson Lab | Choroid plexus tissue | Volume: 60 x 60 x 60 $\mu\text{m}$<br>Resolution: 8 x 8 x 8 nM voxel size |
| Mouse liver<br>(https://doi.org/10.25378/janelia.16913047.v2) | Mouse (P8), C57BL/6J from Jackson Lab | Liver tissue | Volume: 102 x 101.1 x 71.5 $\mu\text{m}$<br>Resolution: 8 x 8 x 8 nM voxel size |
| Mouse kidney<br>(https://doi.org/10.25378/janelia.16913035.v2) | Mouse (P8), C57BL/6J from Jackson Lab | Kidney tissue | Volume: 102 x 101.1 x 71.5 $\mu\text{m}$<br>Resolution: 8 x 8 x 8 nM voxel size |
| Mouse thymus<br>(https://doi.org/10.25378/janelia.24085287.v1) | Mouse (P7), C57BL/6J from Jackson Lab | Thymus tissue | Volume: 147 x 116 x 116 $\mu\text{m}$<br>Resolution: 8 x 8 x 8 nM voxel size |
| Mouse pancreas<br>(https://doi.org/10.25378/janelia.23411843.v2) | Mouse (P7), transgenic mouse with tdTomato-neurons, C57BL/6J from Jackson Lab | Pancreas | Volume: 146 x 113 x 115 $\mu\text{m}$<br>Resolution: 8 x 8 x 8 nM voxel size |
| Mouse skin<br>(https://doi.org/10.25378/janelia.24085269.v1) | Mouse (P7), C57BL/6J from Jackson Lab | Skin | Volume: 138 x 114 x 158 $\mu\text{m}$<br>Resolution: 8 x 8 x 8 nM voxel size |
| Mouse Meissner corpuscle<br>(https://doi.org/10.25378/janelia.23969076.v1) | Mouse (P21), C57BL/6J | Meissner Corpuscle | Volume: 37 x 46 x 58 $\mu\text{m}$<br>Resolution: 6 x 6 x 6 nM voxel size |
| Mouse pacinian corpuscle<br>(https://doi.org/10.25378/janelia.23969112.v1) | Mouse (P112), mixed background Plp1-EGFP mouse | Pacinian Corpuscle | Volume: 58 x 63 x 291 $\mu\text{m}$<br>Resolution: 6 x 6 x 6 nM voxel size |
| Mouse hair follicle<br>(https://doi.org/10.25378/janelia.23969061.v1) | Mouse (P21), C57BL/6J | Hair Follicle | Volume: 88 x 96 x 78 $\mu\text{m}$<br>Resolution: 6 x 6 x 6 nM voxel size |
| Human placenta<br>(doi: [10.1016/j.jisci.2022.105453](https://doi.org/10.1016/j.jisci.2022.105453)) | Human placenta from uncomplicated human pregnancy | Placenta | Various volumes and resolutions |

### Cell annotations and density analysis

We performed cellular (EC and MC) and subcellular (centriole and primary cilium) annotations in pial arterioles and venules, penetrating arterioles and ascending venules, and arteriolar and venular end capillaries. In total, 27 arterioles and 40 venules were analyzed: 17 arterioles and 27 venules in the MICrONS dataset, 5 arterioles and 5 venules in CA1, and 5 arterioles and 8 venules in h01. Capillaries analyzed were 1^st^-3^rd^ order branches from both arteriolar and venular ends. In the human (h01) dataset, due to limited number of capillaries directly branching off from arterioles and venules, we categorized capillaries as either arteriolar-end or venous-end capillaries. Capillaries that were not proximal to penetrating arterioles or ascending venules were labeled as high-order capillaries. ECs were identified as those forming the innermost lining of blood vessels, while MCs (smooth muscle cells or pericytes) were located in the adjacent layer, sharing a basement membrane with the endothelium. Cell positions (coordinates) within the datasets were recorded, and these coordinates were used to calculate the distances between annotations. The center of each cell was determined by identifying the middle z-plane of the nucleus. For cell density calculations, surface area was estimated using cylindrical formulas, and the total number of cells annotated per vessel was divided by the corresponding surface area.

Centrioles are mostly surrounded by the Golgi apparatus, which facilitated the identification of centrioles and primary cilia by first locating the Golgi, typically near the nucleus. The area adjacent to the Golgi was examined for cylindrical structures, which are characteristic of centrioles. In some cases, centrioles were not well-stained and exhibited minimal electron density; however, the gaps they created between organelles of the Golgi apparatus were still be recognized.

### Cilia ultrastructure analysis

Cilia ultrastructure was assessed in a little over half of the annotated ciliated MCs cells in the MICrONS volume and all the h01 annotated vascular cells. The cilia origin was evaluated, and cilia were classified pocket cilia, surface cilia, partial pocket cilia, concealed cilia, unclear, no cilium, or ciliary vesicle. The external face of all cilia that emerged from the cell were classified as endothelial-facing, non-endothelial-facing and retained within the basal lamina, neuropil-entering, contacting both an endothelial cell and the neuropil, contacting another cell type such as a fibroblast or macrophage (rare) or unclear. Cilia that passed beyond the basal lamina were assessed to determine if they were ensheathed by an astrocytic endfeet. We also annotated when a pocket or partial-pocket cilium exited the pocket but then at the tip was ensheathed by its own cell membrane. The fractional distributions of cilia ultrastructure were quantified for the penetrating arteriole, the 1^st^-3^rd^ order capillaries and the ascending venule. Comparisons of the arteriolar end to the venule end also include the cilia in white matter and pial vessel cells.

### Statistical analysis

Data were processed in Microsoft Excel and statistical analyses were performed using GraphPad Prism 10 (GraphPad, USA). Data are presented as frequency distributions and mean ± SEM. Data were tested for normality using the Shapiro–Wilk test prior to application of the appropriate parametric or nonparametric statistical test. One-way ANOVA with post hoc analysis or student’s t-test results are represented in the figure legends and reported in the results text. A p < 0.05 was considered statistically significant. Stars denote significant differences; “ns” indicates nonsignificant comparisons. Linear regression and Pearson’s correlation coefficient were used to evaluate relationship between vascular cell number with vessel surface area.

## Results

### Human microvascular endothelial cells have primary cilia

Given the critical role of ECs in NVC ^10,52–56^ and the differences in reports of cilia abundance in the ECs ^21,22,24,27,29,30^, we examined EC ciliation in the brain datasets. We utilized three large volume EM datasets: mouse visual cortex (MICrONS MM^3; hereafter MICrONS; ^47^), mouse hippocampus (CA1 region ^48^) and human cerebral cortex (h01 ^49^). The MICrONS and CA1 datasets generated from perfusion fixed brain tissue, whereas the h01 dataset was derived from post drop-fixed human cortical tissue, which resulted in some structural deformities, including swollen astrocytes, thick basal lamina and compressed vascular lumen (**Sup. Figure 1**). We reconstructed the vasculature from pial and penetrating arterioles through capillaries to ascending and pial venules in all three datasets. Pial arterioles and venules were distinguished based on MC coverage, with arterioles exhibiting a dense layer of MCs (SMCs) and pial venules showing a more dispersed MC arrangement. These pial vessels were traced through the parenchyma to segment the corresponding penetrating and ascending vessels as well as the surrounding capillaries (**Figure 1a-c**). ECs and MCs were identified and annotated based on their proximity to the vessel lumen, as well as their shape and morphology (**Figure 1d**). We performed a comprehensive analysis of primary cilia in all annotated ECs. In the mouse cortex and CA1 regions, we did not observe primary cilia in ECs (**Figure 1e, f**). In contrast, in the human cortex, approximately 18% of ECs exhibited primary cilia (**Figure 1f, g**). These ciliated ECs were most frequently located in arteriolar-end capillaries (∼40%), followed by penetrating arterioles, ascending venules and venous-end capillaries (14.5-19.5%) (**Figure 1h**). Primary cilia on penetrating arteriole ECs were the longest (∼1-2 μm), compared with those on ascending venule and capillary ECs (**Figure 1i**). Together, these findings demonstrate that ECs in the human, but not in the mouse brain, possess primary cilia, highlighting a clear ultrastructural difference between species.

**Figure 1:**
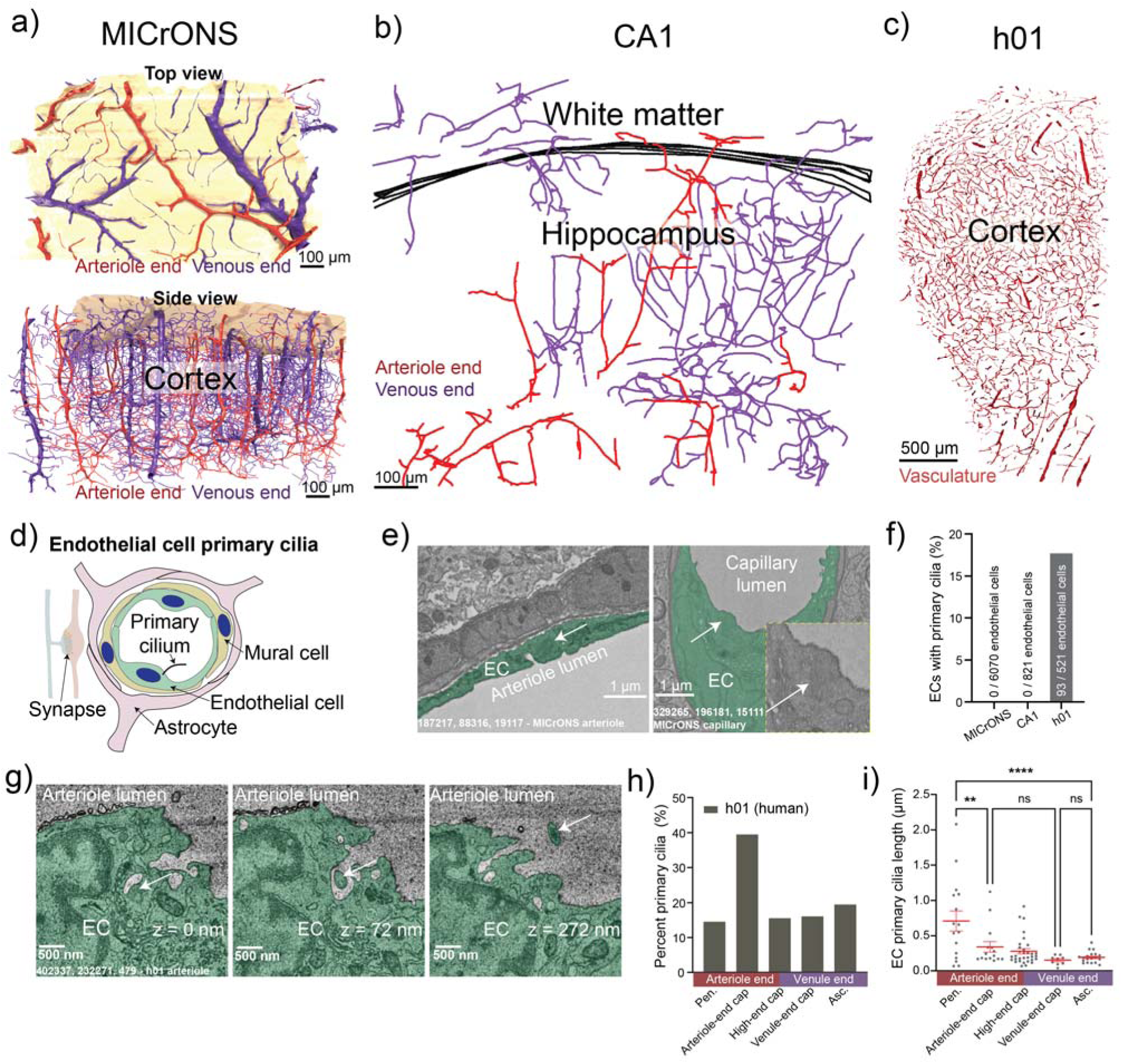
Human endothelial cells have primary cilia, but not mouse. Three-dimensional representations of brain microvasculature from distinct datasets: **(a)** MICrONS (mouse visual cortex) - upper panel shows a top view of the pial vasculature and lower panel shows a side view of the parenchymal vasculature, with the brain surface rendered in beige; **(b)** CA1 (mouse hippocampus CA1 region) - vascular skeletonization, with black lines indicating the boundary between hippocampus and white matter; **(c)** h01 (human temporal cortex) - microvascular architecture reconstructed from human tissue. **(d)** A schematic illustration of the neurovascular unit highlighting primary cilia in ECs. **(e)** Representative MICrONS examples of arteriolar (left) and capillary (right) ECs lacking primary cilia. **(f)** Percent distribution of EC primary cilia in MICrONS, CA1, and h01 datasets. **(g)** Representative h01 example showing a human arteriolar EC primary cilium protruding into the arteriole lumen (left to right, arrows indicate cilium). **(h)** Percent distribution and **(i)** length of EC primary cilia within human cortical vasculature, separated into arteriolar and venous ends. Data shown as mean ± SEM (** p<0.01, **** p<0.0001, one way ANOVA with post-hoc Tukey’s test).

**Supplemental Figure 1:**
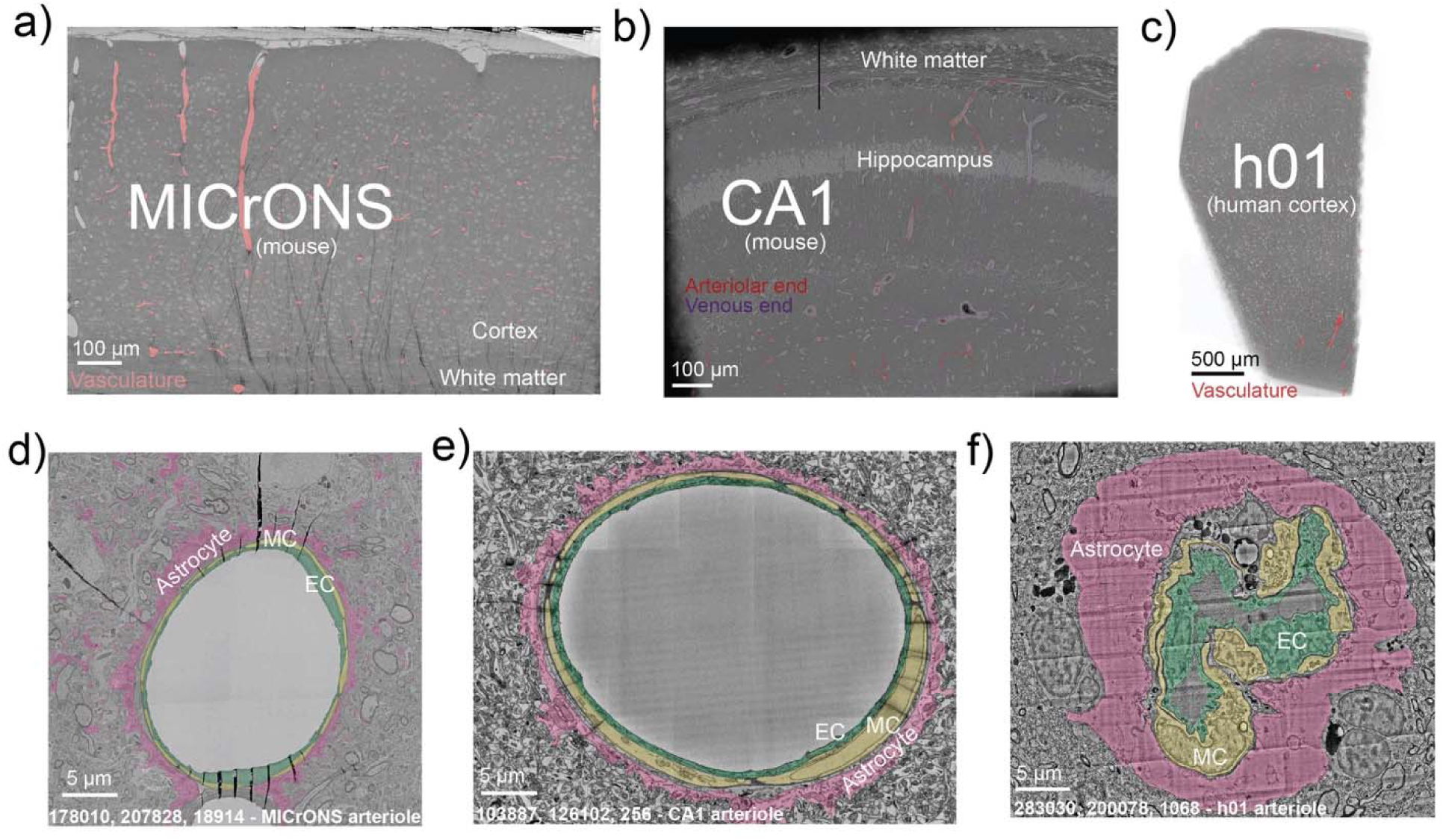
Astrocyte swelling in the h01 dataset. **a-c)** Electron micrographs with vasculature indicated from MICrONS (mouse visual cortex), CA1 (mouse hippocampus CA1 region), and h01 (human temporal cortex) datasets used for segmenting and annotating microvasculature and vascular cells. **(d-f)** Examples of arterioles showing ECs (green), MCs (yellow) and astrocytes (pink) from (d) MICrONS, (e) CA1 and (f) h01, respectively. Note the slender astrocytic endfeet in perfusion-fixed mouse samples (MICrONS and CA1) compared to the swollen endfeet in the drop-fixed human sample (h01), which compress the vasculature.

### Brain mural cells have primary cilia

Primary cilia in MC are described only in a few studies. Biscoe et al. reported primary cilia in pericyte of the carotid body, where cilia shown to project toward the vascular lumen in blood vessels ^37^. Another study identified primary cilia in approximately 80% of brain pericytes using immunofluorescence techniques ^39^. However, these studies did not examine differences in pericyte primary cilia across the microvascular network, nor did they compare their presence or ultrastructure across species and tissues. This left a critical gap in our knowledge about the distribution and morphology of MC primary cilia within the brain microvasculature (arterioles, capillaries and venules). To address this gap, we systematically evaluated the presence and ultrastructure of MC primary cilia across the cerebral microvasculature and between species (**Figure 1a-c, 2a**). Unlike ECs, we identified MC with primary cilia in both mouse and human brains, oriented either towards ECs or astrocytes. Figure 2b shows an example of a MC primary cilium oriented towards an EC in the MICrONS dataset. Approximately 40-50% of MCs were ciliated in both species (Figure 2c), with a gradient in primary cilia prevalence across the microvasculature (Figure 2d). In the MICrONS datasets, 83% of pial venular and 92% of ascending venular MCs possessed primary cilia, whereas only 3% of pial arteriolar and <1% penetrating arteriolar MCs displayed them (Figure 2e). A similar trend was observed in CA1 datasets, 92% ascending venular MCs possessed primary cilia, whereas none of penetrating arteriolar MCs were ciliated (Figure 2f). In both the MICrONS and CA1 datasets, primary cilia distribution also differed between arterial- and venous-end capillaries, depending on their proximity to penetrating vessels. MCs (pericytes) associated with capillaries near venules exhibited the highest frequency and greatest length of primary cilia (Figure 2d-f). In the human brain (h01), we again observed frequent primary cilia on ascending venules (∼62%) and capillaries but detected only a single primary cilium on a penetrating arteriole MC (**Figure 2d, g**), despite arterioles containing a greater number of MCs (SMCs). Mouse MC cilia were significantly longer (∼5 μm) (Figure 2e) compared to human MC cilia (∼2 μm) (Figure 2i). Given the observed gradient in both cilia distribution and length, we extended our analysis to MCs in other brain and peripheral tissues (Figure 2h-j). Interestingly, ciliated MCs (pericytes) were also identified in macaque cortex, mouse white matter, heart, choroid plexus, liver, placenta, kidney, pancreas, thymus, skin, hair follicle, meissner and pacinian corpuscles of the dermis (Figure 2i-j and Sup. Figure 2), where cilia lengths averaged approximately 2 μm. In summary, these data reveal a gradient distribution of primary cilia in MCs across the brain vasculature, with the highest prevalence in the venous end of the microvasculature.

**Figure 2:**
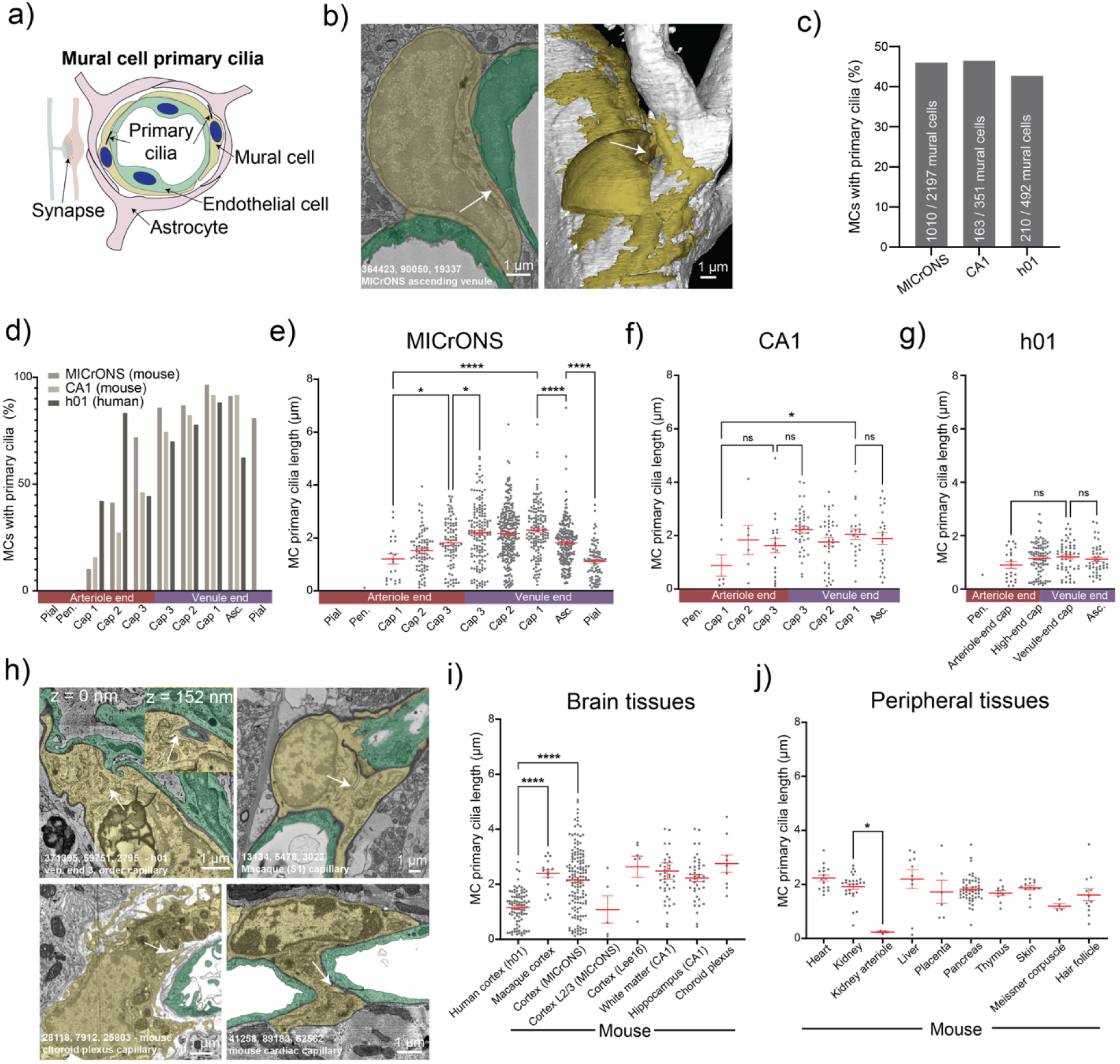
Mural cell primary cilia prevalence and distribution. **a)** A schematic illustration of the neurovascular unit highlighting primary cilia on mural cells. **(b)** Example from MICrONS dataset showing: *Left*: a MC primary cilium (orange, indicated by arrow) aligned next to an EC (green), *right*: the 3D structure of the MC (yellow) and the venous-end microvasculature (white). **(c)** Percent distribution of primary cilia in MCs across the three datasets. **(d)** Percent distribution of primary cilia in MCs after microvascular stratification into pial and parenchymal vessels, including 1-3^rd^ order capillaries from the MICrONS (mouse visual cortex), CA1 (mouse hippocampus), and h01 (human cortex) datasets. Note that neither CA1 nor h01 contain any pial vessels. **(e-g)** Lengths of MC primary cilia in pial and parenchymal vessels, including 1-3^rd^ order capillaries MCs from **(e)** MICrONS, **(f)** CA1 and **(g)** h01 datasets. **(h)** Representative examples of pericyte primary cilia in human cortex (**top left**), macaque cortex (**top right**), mouse choroid plexus (**bottom left**) and mouse heart (**bottom right**). **(i-j)** Length of MC primary cilia from **(i)** other brain regions and **(j)** peripheral tissues. Data shown as mean ± SEM (* p<0.05, ** p<0.01, **** p<0.0001, ns: not significant, one-way ANOVA with post-hoc Tukey’s test).

**Supplemental Figure 2:**
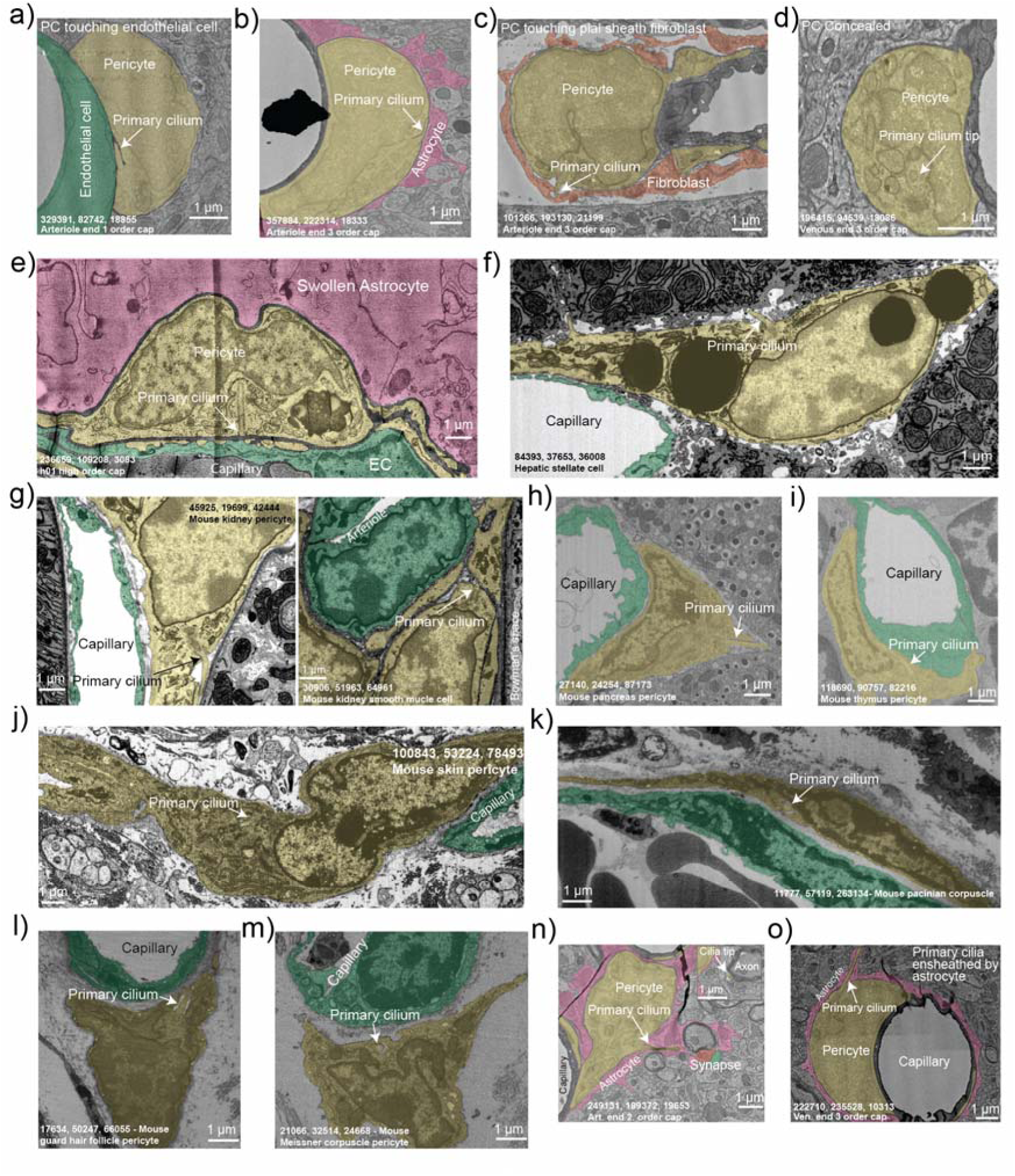
Examples of mural cell primary cilia in various tissues. **(a-c)** MC primary cilia in contact with **(a)** an EC, **(b)** an astrocyte, and **(c)** a fibroblast. **(d)** Example of a MC primary cilium that remains entirely within the cytosol. **(e)** Human MC primary cilium with a very elongated mother centriole, surrounded by a swollen astrocyte. **(f)** Mouse hepatic stellate cell in the liver with a primary cilium. **g)** Kidney: primary cilia on a pericyte (left) and a smooth muscle cell (right). **h)** Mouse pancreas pericyte with a primary cilium. **i)** Mouse thymus pericyte with a primary cilium. **j)** Mouse skin pericyte with a primary cilium. **k)** Mouse Pacinian pericyte with a primary cilium. **l)** Mouse guard hair follicle pericyte with a primary cilium. **(m)** Mouse Meissner corpuscle pericyte with a primary cilium. **(n)** MC primary cilium penetrating between astrocyte endfeet to come in direct contact with the neuropil (see inset from a different layer showing the primary cilium tip). **(o)** MC primary cilium ensheathed by astrocyte endfeet. Arrows indicate the locations of the primary cilia.

### Species-specific differences in primary cilia ultrastructure, orientation, and the surrounding extracellular environment

During our examination of centrioles and primary cilia, several ultrastructural features were observed. Many non-ciliated centrioles were associated with a membrane. In these cells, distal appendages of the mother centriole were either docked directly at the cell surface or tethered to internal membranes such as tubules or vesicles (Figure 3a). These structures may represent cilia assembly or disassembly intermediates ^42,57^.

**Figure 3:**
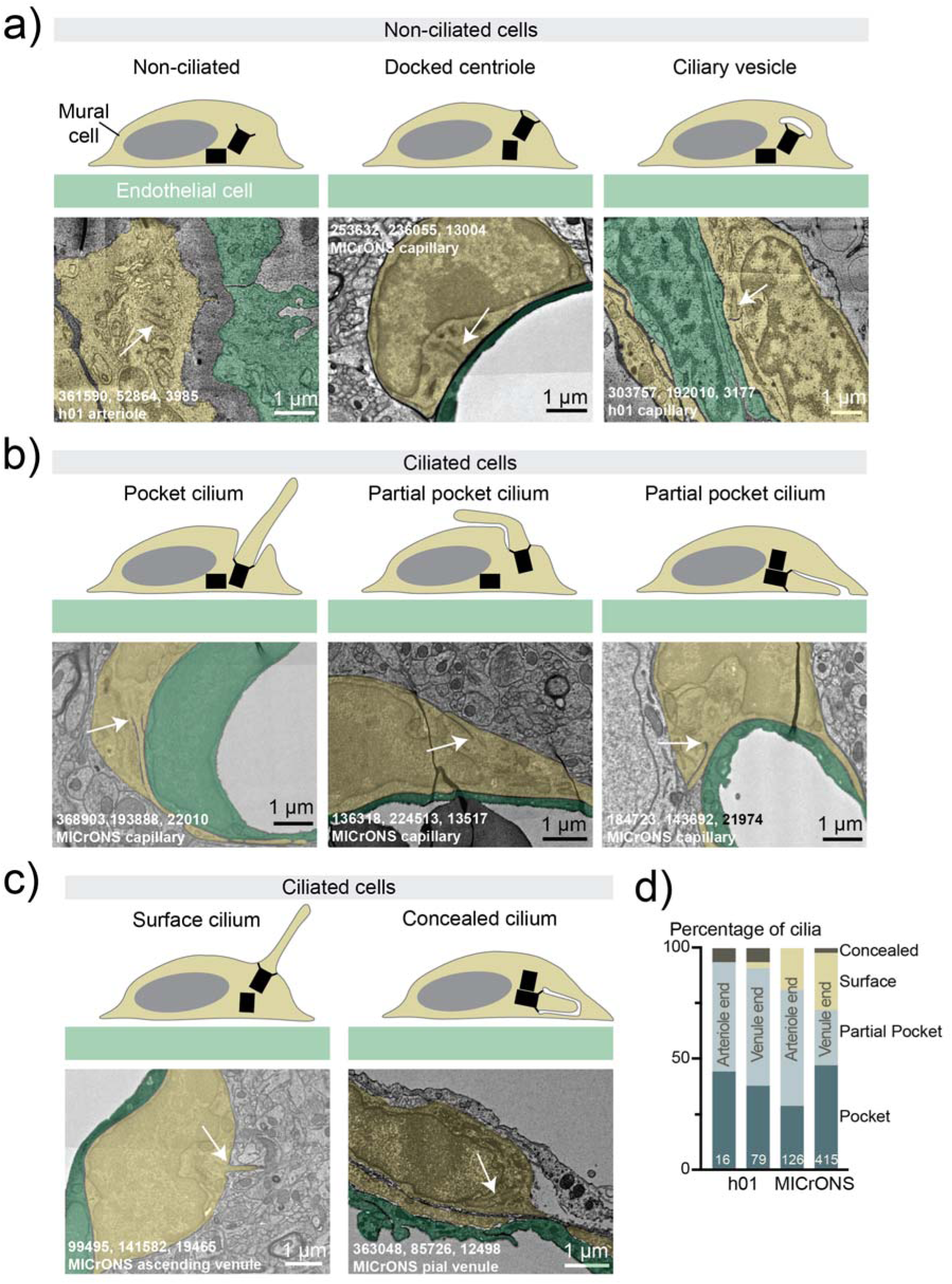
Heterogeneity in the mural cell ciliogenesis. Schematic illustration (top) and examples (bottom) of different forms of **(a)** non-ciliated and **(b-c)** ciliated cells present in MICrONS and h01 datasets. White arrows indicate the centrioles or base of the primary cilia. **(d)** Comparison of the percentage of cilia facing endothelial cells versus the parenchyma in MICrONS and h01 datasets.

In ciliated MCs from the MICrONS and h01 datasets, we assessed the structure of the ciliary bases. Typically, cilia form either with a ciliary pocket—where the base is recessed into the cytoplasm—or at the cell surface, depending on the biogenesis pathway ^15^. Surface cilia grow from plasma membrane–docked centrioles, whereas pocket cilia begin within the cytoplasm with their nascent axoneme enveloped by a membrane that later fuses with the plasma membrane. Because the centriole remains distant from the cell surface, the membrane enveloping the nascent cilium becomes the invaginating membrane of the ciliary pocket ^17,58^. While examples of pocket cilia were present across all datasets, surface cilia were rare in the human dataset (**Figure 3b, d**). We also identified a small proportion of cells with concealed cilia—internal cilia that have not reached the plasma membrane—previously described in neurons and glia (**Figure 3c, d;** ^41,42^). In both h01 and MICrONS datasets, a large fraction of MCs (25-50%) had centrioles with a subset of distal appendages anchored at the plasma membrane and the rest anchored to an invaginating membrane, resulting in a partial pocket (**Figure 3b, d**). Photoreceptors are the only other cells we are aware of that have a partial pocket. The observed non-uniformity of cilia origins may indicate differences in the ciliogenesis.

In addition to structural analysis of the cilium itself, volume EM enabled examination of the surrounding environment. For each cilium, we annotated whether it projected toward parenchyma or endothelial cells. There was a substantial difference between MC cilia in the MICrONS datasets and h01: approximately 75% were oriented toward parenchyma in mouse cortex whereas nearly all cilia emerged adjacent to an EC in human cortex (**Figure 4a-c**). We wondered if this difference in orientation could be related to the differences in frequency of surface, pocket, and partial pocket cilia. Upon separate analysis of endothelial- and parenchyma-facing cilia in the MICrONS dataset, we found that endothelial-facing cilia were rarely surface cilia (**Figure 4d,e**). Thus, the scarcity of surface cilia observed in the human dataset (**Figure 3d**) may be explained by the fact that endothelial-facing cilia are less likely to grow directly from the cell surface, possibly due to spatial constraints.

**Figure 4:**
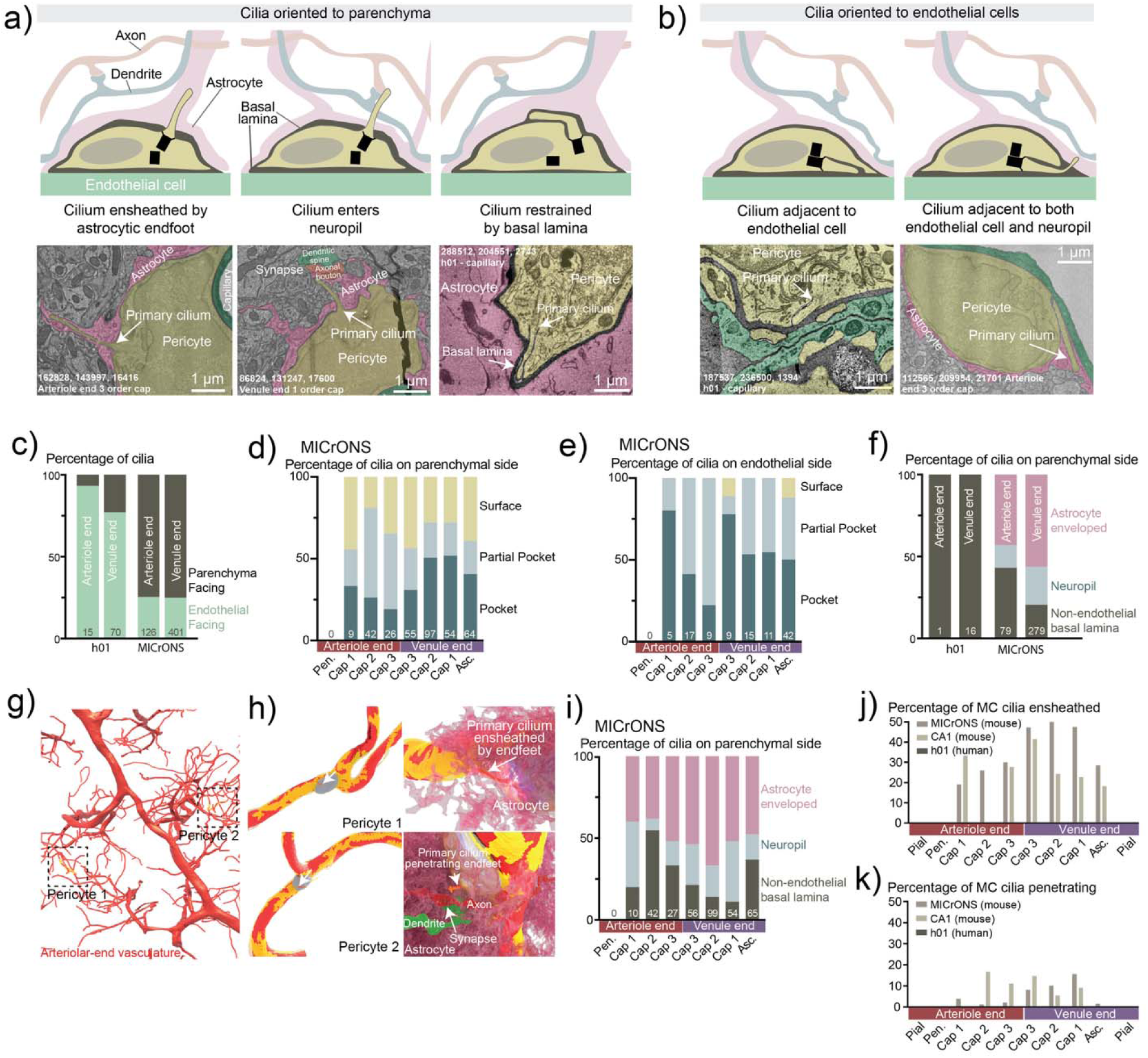
Mural cell primary cilia location and orientation highlighting interspecies differences. **(a-b)** Schematic representation of MC primary cilium positions relative to surrounding structures. **(a)** Examples of MC primary cilium (top: schematic; bottom: EM reconstructions) (left) ensheathed by astrocytic endfeet, (middle) entering the neuropil, and (right) constrained by the basal lamina. **(b)** Example of MC primary cilium adjacent to both an EC and an astrocyte. (**c**) The distribution of analyzed cilia facing either the endothelial cell or the parenchyma is graphed for cells of the arteriole or venule sides of the vasculature for the h01 and MICrONs datasets. (**d-e**) The fraction of analyzed cilia in MICrONs with surface, pocket or partial pocket origins were graphed for the indicated vessel types for **(d)** endothelial facing cilia or **(e)** parenchyma facing cilia. (**f**) The parenchyma facing cilia were further classified and the percentage of cilia constrained by the basal lamina, entering the neuropil, or entering the neuropil and enveloped by an astrocyte endfoot are graphed for the arteriole and venule ends of the h01 and MICrONs datasets. **(g)** Overview of arteriolar-end microvasculature indicating locations of the two pericyte segmentations shown in **h** (dotted box). **(h)** Segmentations of arteriolar-end capillary pericytes (yellow) with primary cilia (orange) and nuclei (gray). *Left panels*: segmented pericytes. *Right panels*: interactions of their primary cilia with neural cells. *Top*: example of a pericyte primary cilium ensheathed by astrocyte endfeet (pink). *Bottom*: example of a pericyte primary cilium penetrating between the astrocyte endfeet (pink) to come in close contact with an axonal bouton (red) and a dendritic spine (green) forming a synapse. (**i**) The percentage of parenchyma-facing cilia from MICrONs constrained by the basal lamina, entering the neuropil or entering the neuropil and enveloped by an astrocyte endfoot is graphed for each indicated subset of vessels. **(j-k)** Percentage of MC primary cilia **(j)** ensheathed by astrocyte endfeet and **(k)** penetrating between astrocyte endfeet on pial and parenchymal vessels, including 1-3^rd^ order capillaries.

Further analysis of parenchyma-oriented cilia revealed additional interspecies differences (**Figure 4f**). Many MC cilia in the MICrONS dataset pass through the basal lamina and ensheathed by astrocyte endfeet (**Figure 4a, g, Sup. Figure 3, Sup. Video 1**), and once in the mouse neuropil, some cilia wrapped between neural processes, a few passing close to synapses (**Figure 4h, Sup. Figure 3, Sup. Video 2**). In the pia, cilia that passed the basal lamina could be found adjacent to fibroblasts. Most strikingly, the majority of cilia were ensheathed, partially or completely, by an astrocytic endfeet, with their tips indenting the astrocytic plasma membrane and extending deep into the astrocyte process (**Figure 4a, g-j**). In a small subset of capilary pericyte primary cilia in the mouse datasets, we found cilia penetrating into the neuropil that were not ensheathed by astrocyte endfeet (**Figure 4k**). In the human dataset where cilia were shorter (∼1 µm), cilia were confined by a thick basal lamina which prevented entry into the neuropil (**Figure 4a, f**).

The close association of the cilium with the astrocyte might shield a cilium from outside signaling molecules and/or enable targeted signaling from the astrocyte to the MC cilium. To quantify this phenomenon, we assessed the frequency of astrocyte envelopment across the microvasculature in the MICrONs dataset; in regions with ciliated MCs, 38-60% of cilia were ensheathed (**Figure 4i, j, Sup. Figure 3, Sup. Video 3**). Similar analyses of the mouse CA1 dataset revealed that the frequency of astrocyte ensheathment was similar across vasculature types and always exceeded the fraction of non-ensheathed cilia (**Figure 4j**). The intimate relationship of MC cilia with astrocytes is suggestive of an important physiological role, however, the finding that some mouse MC cilia and the majority of human MC cilia are adjacent to ECs suggests that different MC cilia could be positioned to encounter signals released from multiple sources.

**Supplemental Figure 3:**
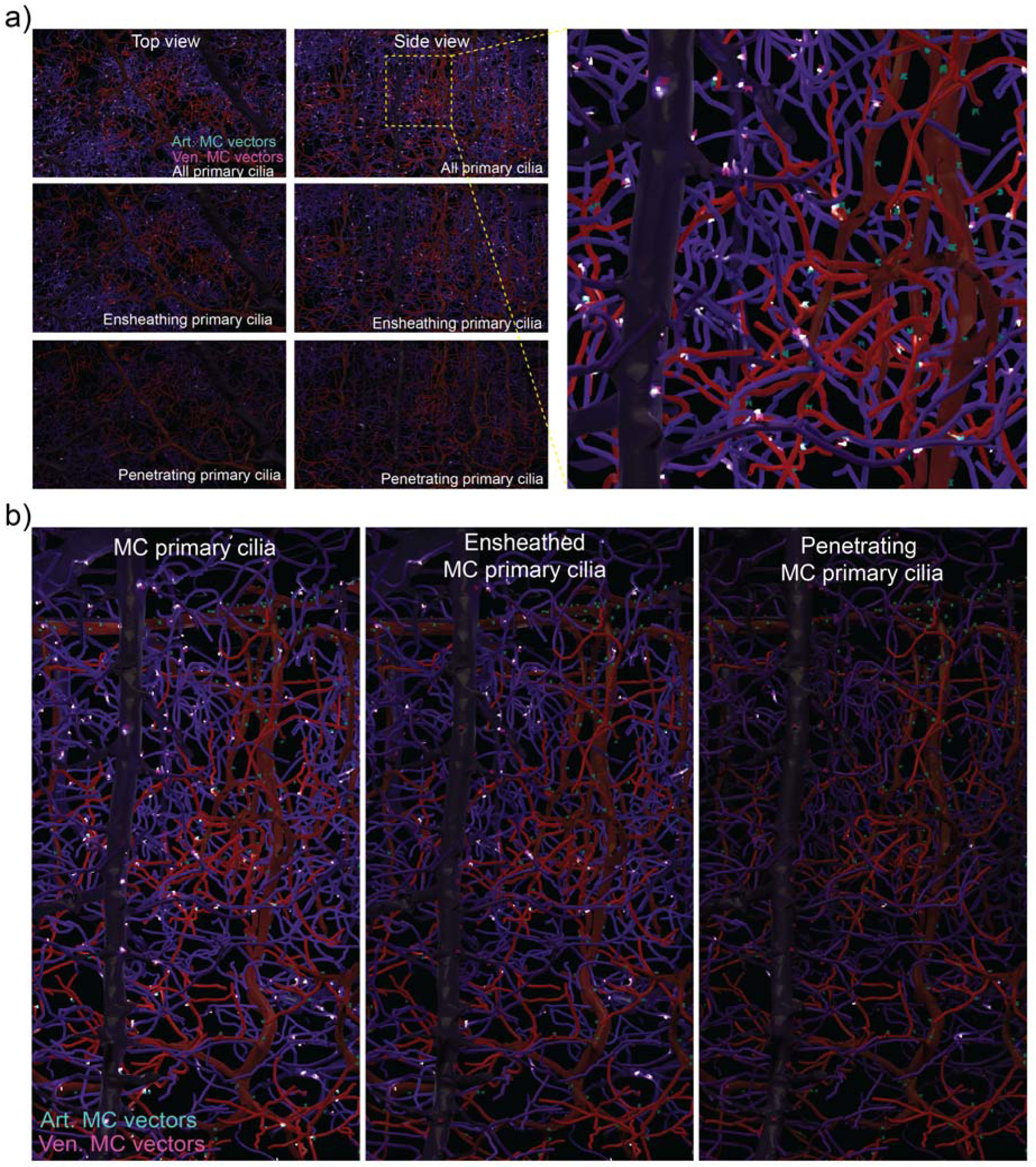
Locations of mural cell primary cilia in the MICrONS dataset. To illustrate the location of MC primary cilia in the MICrONS dataset, we made primary cilia vectors from the base to the tip of the cilia, turned down the light and let the vectors glow. **(a)** *Top panels*: Shows all of the primary cilia in the MICrONS dataset from a top view (left) and a side view (right). The yellow dashed lines indicate the enlarged side view to the right, where mural cell nucleus-to-centriole vectors are visible and primary cilia are glowing. *Middle panels*: Shows only the primary cilia vectors of MC primary cilia that are ensheathed by astrocytes. *Lower panels*: Shows only the primary cilia vectors of MC primary cilia that penetrate between astrocyte endfeet to be in direct contact with neuropil. **(b)** Three enlargements of the side views from **a**.

### Endothelial and mural cell density across the brain microvasculature

Large-volume EM datasets allowed us to quantify vascular cell (EC and MC) densities with high precision. Vascular cells form and maintain the BBB and play key roles in NVC through intercellular communication ^5,59^. EC density varies with local shear stress and the mechanical demands placed on the vessel wall, whereas density of MCs may reflect their capacity to regulate vascular tone and blood flow ^5^. However, their cellular densities along the vascular continuum remain poorly identified. Although It is generally recognized that MCs are more densely distributed at the arteriolar end compared to the venular end ^1,7^, however a systematic quantification of EC and MC densities across the microvasculature is warranted. While previous reports have identified vascular length in the mouse cortex (∼0.88 m/mm^3^) ^60^, macaque (∼0.4-0.5 m/mm^3^) ^61^ and human cortex (∼0.2-0.5 m/mm^3^) ^62,63^, these studies did not measure the density of vascular cells. The high-resolution large-volume EM datasets provided an opportunity to revisit this description and perform a comprehensive analysis of total vascular cell density per vessel lumen surface area in each dataset, comparing the arteriolar and venous vasculature (**Figure 5a**).

**Figure 5:**
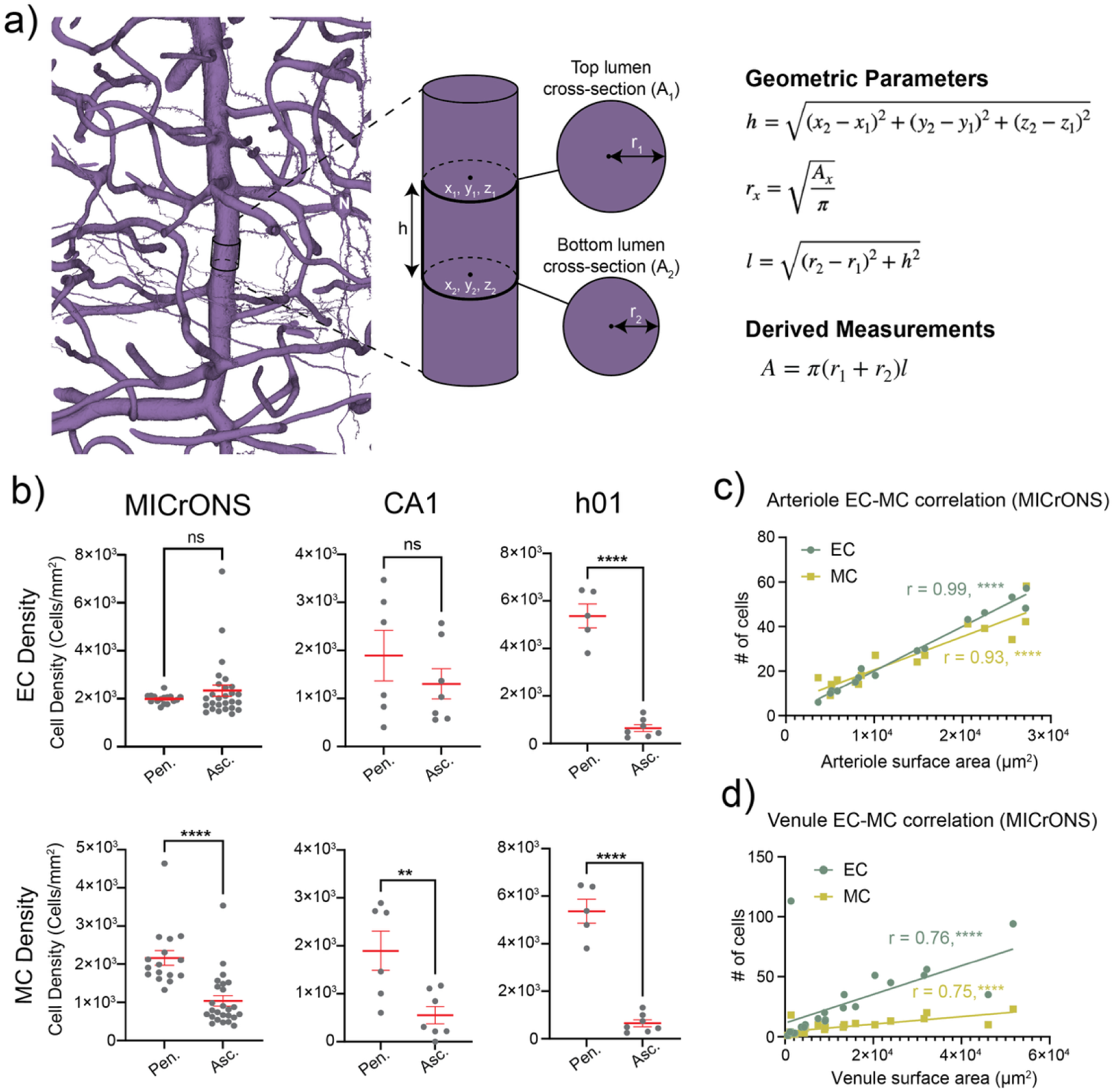
Mural cells have higher density in the arteriolar end, while endothelial cells are similar in both ends. **a)** Illustration with approach used for calculating surface area**. (b)** Vascular cell densities of arterioles and venules (as cells per microvascular lumen surface area). *Top panels*: EC densities and *Bottom panels*: MC densities in the MICrONS (mouse cortex), CA1 (mouse hippocampus) and h01 (human cortex) datasets. Each point represents a single blood vessel; significance was tested with an unpaired t-test (ns: not significant, ** p<0.01, **** p<0.0001). **(c)** Correlation of cell number and arteriole surface area for ECs and MCs. Pearson’s correlation (r > 0.9) shows a strong positive relationship between vascular cell density with arteriolar surface area. **** p<0.001. **(d)** Correlation of cell number and venule surface area for ECs and MCs. Pearson’s correlation (r ∼ 0.75) shows a positive relationship between vascular cell density with venous surface area. **p<0.01, ****p<0.0001, ns: not significant.

We first compared ECs densities in parenchymal (**Figure 5b, top left**) and pial (**Sup. Figure 4a-c**) vessels. In mouse cortex (MICrONS), all pial arterioles and venules had comparable EC densities of ∼2500-2900 ECs per square millimeter of vessel surface area. Similarly, parenchymal arterioles and venules also had comparable EC density (∼1900-2400 cells/mm^2^). Our analysis of hippocampus/CA1 vasculature also showed comparable EC densities in parenchymal arterioles and venules (∼1300-1900 ECs/mm^2^), with no significant differences (**Figure 5b, top center**). In contrast, human parenchymal arterioles had approximately twice the EC density (2818 ± 423 cells/mm^2^) compared to parenchymal venules (1163 ± 187 cells/ mm^2^) (**Figure 5b, top right**). Differences in fixation methods between the mouse and human tissue could account for the observed difference between species because swelling in the h01 could have altered the vessel surface area. In mouse, however, EC densities were generally comparable across arteriolar and venous.

We next analyzed MC density and found that pial and parenchymal arterioles had significantly higher cell densities compared to pial and parenchymal venules in both mouse and human brain (**Figure 5b, bottom**). In the Mouse cortex (MICrONS), pial and parenchymal arterioles contained approximately 2613 ± 254 cells/mm^2^ and 2164 ± 197 cells/mm^2^, respectively, which were more than double the density observed in pial (625 ± 57 cells/mm^2^) and parenchymal venules (1040 ± 138 cells/mm^2^) (**Figure 5b bottom left and Sup. Figure 4c**). In mouse hippocampus/CA1, MC density was 1896 ± 407 cells/mm^2^ in arterioles and 552 ± 180 cells/mm^2^ in venules (**Figure 5b, bottom center**). In human cortex, parenchymal arterioles had ∼5400 cells/mm^2^, which was 9-10-fold higher than parenchymal venules (∼650 cells/mm^2^, **Figure 5b, bottom right**). In summary, MC densities were consistently higher in arterioles than in venules.

Given the similar EC density between mouse arterioles and venules, we next asked whether cell distribution depends on the vessel diameter. To address this, we examined the correlation between cell number and vessel surface areas of individual vessels. In the mouse cortex (MICrONS), the numbers of vascular cells were positively correlated with surface area (correlation coefficient r > 0.9 for the arteriolar end and r ≈ 0.75 for the venous end) (**Figure 5c-d and Sup. Figure 4d**). In contrast, no significant correlation was observed between cell number and vessel surface area in the human cortex (h01) or the mouse CA1 region (**Sup. Figure 4e and f**), despite comparable arteriolar and venular surface areas (**Sup. Figure 4a**). The differences in cell number-surface area correlation between different datasets may attribute to variations in dataset geometry and tissue quality. The MICrONS dataset is substantially larger than the CA1 dataset, while the h01 dataset is of comparable volume to MICrONS but differs markedly in shape –relatively flat rather than cubic. This flat geometry limits the ability to follow vessels continuously, as vascular segments enter and exit the volume over short distances, whereas the MICrONS dataset allow for more complete vessel reconstructions. Furthermore, the h01 dataset exhibits pronounced astrocyte swelling, which may distort vessel lumen surface area measurements and thereby obscure true relations.

Overall, we found that EC density was comparable between arterioles and venules in both the mouse cortex and CA1 regions, suggesting that differences in BBB properties could not be attributed to variations in EC densities. This appears less clear in the human cortex, where variability in tissue preservation may confound interpretation. In both mouse and human brains, venules contained mural cells, but at lower densities than arterioles.

**Supplemental Figure 4:**
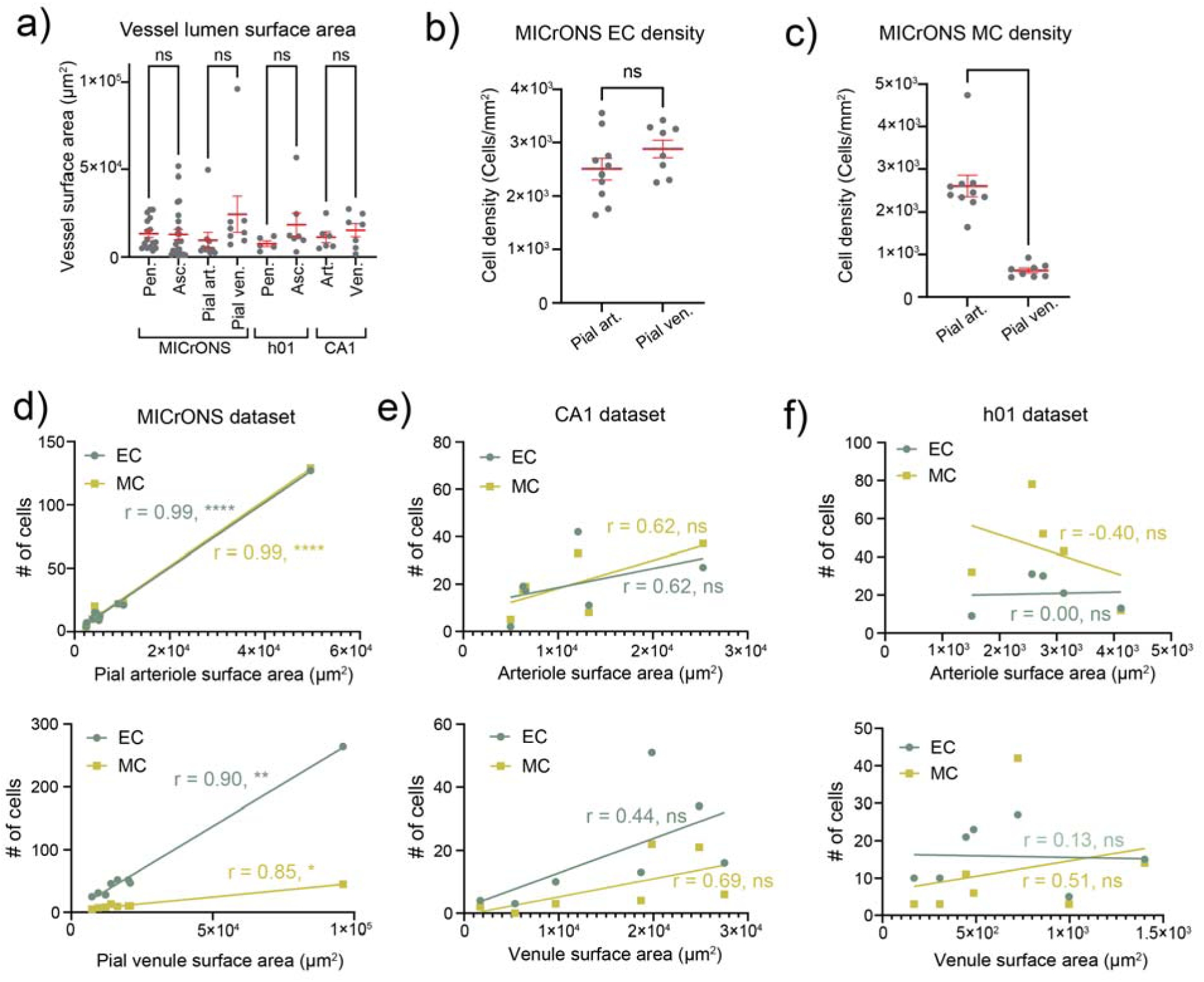
Mural cell and endothelial cell density and surface area. **a)** Mean vascular surface area of arterioles and venules in the MICrONS, h01 and CA1 datasets. **(b)** EC and **(c)** MC cell density as cells per surface area in pial arterioles and venules in the MICrONS dataset. **(d)** *Top panel:* Correlation of cell number and arteriole surface area for ECs and MCs in the MICrONS mouse cortex dataset. Pearson’s correlation (r > 0.9) shows a strong positive relationship between vascular cell density with arteriolar surface area. *Bottom panel:* Correlation of cell number and venule surface area for ECs and MCs. Pearson’s correlation (r ∼ 0.85) shows a positive relationship between vascular cell density with venous surface area. **(e)** *Top panel:* Correlation of cell number and arteriole surface area for ECs and MCs in the CA1 mouse hippocampus dataset. Pearson’s correlation (r = 0.44 (ECs) and 0.69 (MCs)) shows a no significant relationship between vascular cell density with arteriolar surface area. *Bottom panel:* Correlation of cell number and venule surface area for ECs and MCs. Pearson’s correlation (r = 0.62) shows a no significant relationship between vascular cell density with venous surface area. **(f)** *Top panel:* Correlation of cell number and arteriole surface area for ECs and MCs in the h01 dataset. Pearson’s correlation (r = 0.00 (ECs) and −0.40 (MCs)) shows a no significant relationship between vascular cell density with arteriolar surface area. *Bottom panel:* Correlation of cell number and venule surface area for ECs and MCs. Pearson’s correlation (r = 0.13 (ECs) and 0.51 (MCs)) shows a no significant relationship between vascular cell density with venous surface area. * p<0.05, ** p<0.01, **** p<0.0001, ns: not significant.

### Endothelial cells are polarized against blood flow direction

In addition to their role in primary cilium formation, centrioles are essential for organizing microtubules and establishing planer cell polarity (PCP), a coordinated alignment of cells within a 2D plane, where centrioles are typically positioned in the leading edge of the PCP axis ^64–66^. The PCP orientation is strongly influenced by mechanical forces, which could include shear stress generated by blood flow. ECs experience varying degree of shear stress depending on their vascular location: arteriolar ECs are exposed to the highest shear stress associated with high-velocity blood flow, which progressively decreases as blood passes through capillaries and into venules ^67,68^. Despite this well-established gradient, the role of centrioles in EC polarization - and how shear stress variations across the microvasculature influence this process - remains poorly understood. To investigate this, we inferred the CBF direction in all datasets based on vessel identity: pial arterioles conduct blood from larger to smaller surface vessels, pial venules from smaller to larger surface vessels, penetrating arterioles carry blood from the cortical surface into the parenchyma (top-to-bottom), and ascending venules return blood from the parenchyma to the surface (bottom-to-top) (**Figure 6a-b**). We visualized EC nucleus-to-centriole orientations as vectors extending from the center of the nucleus (broad end) to a point between the centrioles (pointed end) (**Sup. Video 4**). We first examined the vector orientation and then the vector lengths. In all three datasets, these vectors consistently aligned opposite to the direction of CBF in both arterioles and venules (**Figure 6a-e**). Notably, across the pial arteriole anastomosis, these vector orientations were reversed, reflecting the opposing CBF directions on either side of the junction. In contrast, vector orientations in pial venules were more heterogenous, which could be attributed to the lower blood flow velocities and reduced shear stress (**Figure 6c and Sup. Figure 5**). In both human and mouse cortex as well as the mouse CA1 region, 94-98% of arteriolar and 82-88% venular EC vectors were oriented opposite to the direction of the CBF (**Figure 6e**). Similarly, 85-92% of capillary EC vectors exhibited the same orientation (**Figure 6f**). These findings strongly suggest that the position of centrioles relative to the nucleus serves as a robust marker of CBF direction and shear stress, even in the vessels with relatively low shear stress and even in fixed tissue.

**Figure 6:**
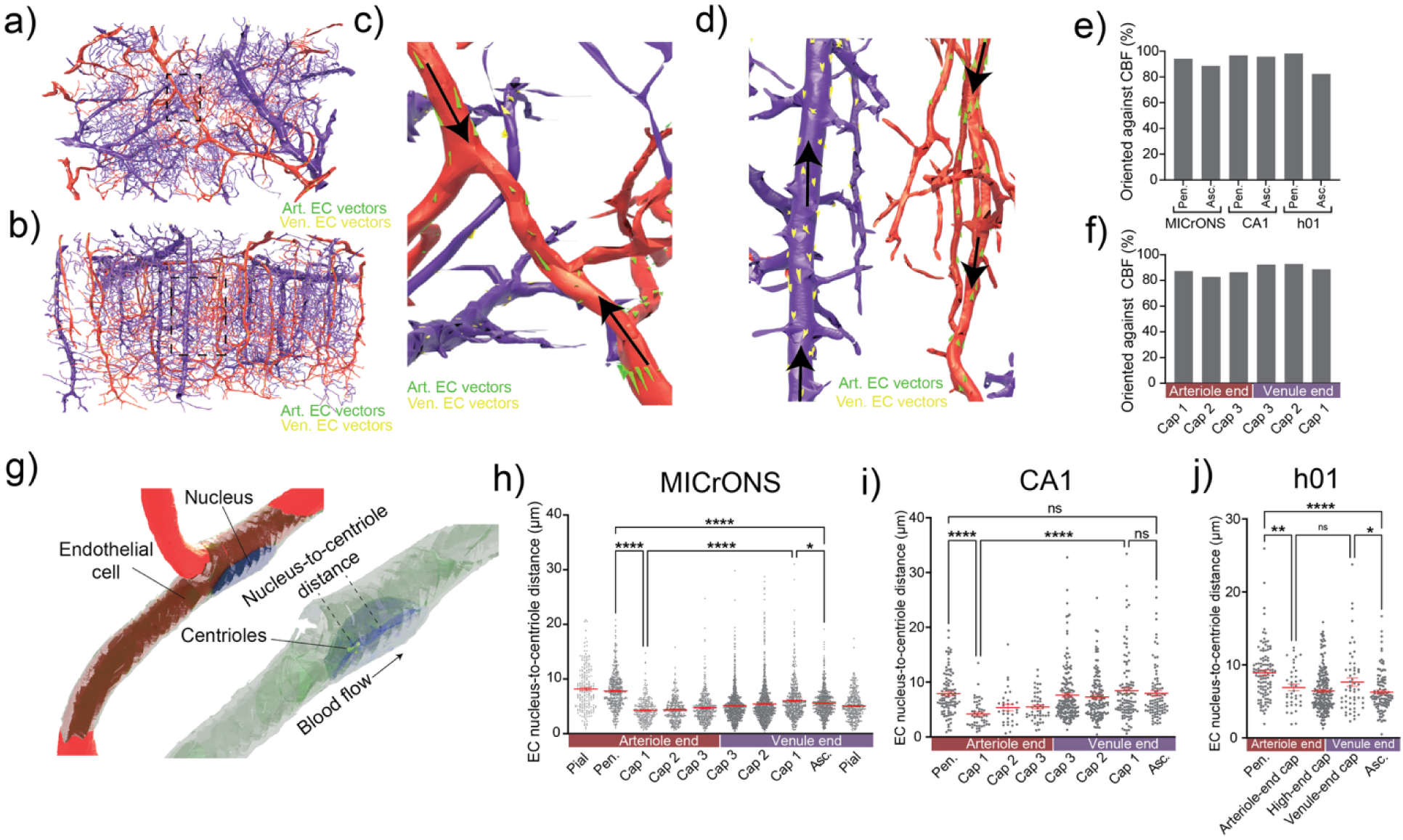
Endothelial cells are polarized against the cerebral blood flow. The MICrONS cortical microvasculature seen from **(a)** top and **(b)** side-view showing arterial venous-end microvasculature with vectors representing the EC nucleus-to-centriole orientations. Dashed black box indicates a subset of vessels that are enlarged in **c-d**. **(c)** Enlargement of an arteriolar anastomosis from the top panel in **a**. Arteriolar nucleus-to-centriole vectors (green) reverse direction across the anastomosis (center of image). **(d)** Arterioles and surrounding capillaries with corresponding vectors (right panel, green vectors) and venules and associated capillaries with corresponding vectors (left panel, yellow vectors). **(e-f)** Percentage of vectors oriented opposite the direction of CBF in **(e)** penetrating arterioles and ascending venules and **(f)** arteriolar and venous end capillaries in MICrONS, CA1 and h01 datasets. **(g)** An example of a segmented endothelial cell showing nucleus and centrioles which were used to create vectors to determine their orientation and calculate nucleus-to-centriole distance. **(h-j)** Nucleus-to-centriole distances in ECs in **(h)** MICrONS, **(i)** CA1 and **(j)** h01. Data are presented as mean ± SEM (*p<0.05, ** p<0.01, **** p<0.0001, One way ANOVA with post-hoc Tukey’s test).

In addition to assessing orientation, the nucleus-to-centriole vectors also provide a measure of the spatial distance between the nucleus and centrioles. This distance reflects cellular morphology, with longer distances indicating a more elongated cell shape. To evaluate morphological differences between ECs and MCs across the vasculature, we quantified nucleus-to-centriole distances in both cell types. **Figure 6g** shows an example of segmented ECs with nucleus-to-centriole vector. On average, these distances were significantly shorter in MCs (∼2-5 µm, **Sup Figure 6**) compared to ECs (∼4-9 µm, **Figure 6h-j**), consistent with the elongated morphology characteristic of ECs (**Figure 6g and Sup. Figure 5**). Among ECs, penetrating arteriolar cells in both mouse and human cortex exhibited the longest nucleus-to-centriole distance (**Figure 6h, j**), suggesting a more elongated morphology in these regions. In contrast, no significant differences were observed between arteriolar and venular ends in the mouse CA1 region (**Figure 6i**). Across all three datasets, nucleus-to-centriole distance consistently decreased from arterioles to capillaries (**Figure 6h-j**), reflecting morphological adaptations to local hemodynamic changes. Notably, capillary ECs displayed a progressive increase in nucleus-to-centriole distance from the arteriolar end to the venous end in both the MICrONS and CA1 datasets, revealing spatial variation in EC morphology across the capillary bed (**Figure 6h-i**).

We next asked whether the lower density of MCs on the venous end (**Figure 5**) results in a more elongated morphology to compensate for the reduced coverage and maintain an arteriolar-equivalent surface area. However, aside for the human dataset (h01), we found no significant differences in nucleus-to-centriole distances between arteriolar and venular MCs (**Sup. Figure 6**), suggesting that venular MCs preserve their cell shape and architecture despite covering a comparable surface area. Altogether, these findings highlight distinct variations in cellular morphology across the vascular tree in both mouse and human brains.

**Supplemental Figure 5:**
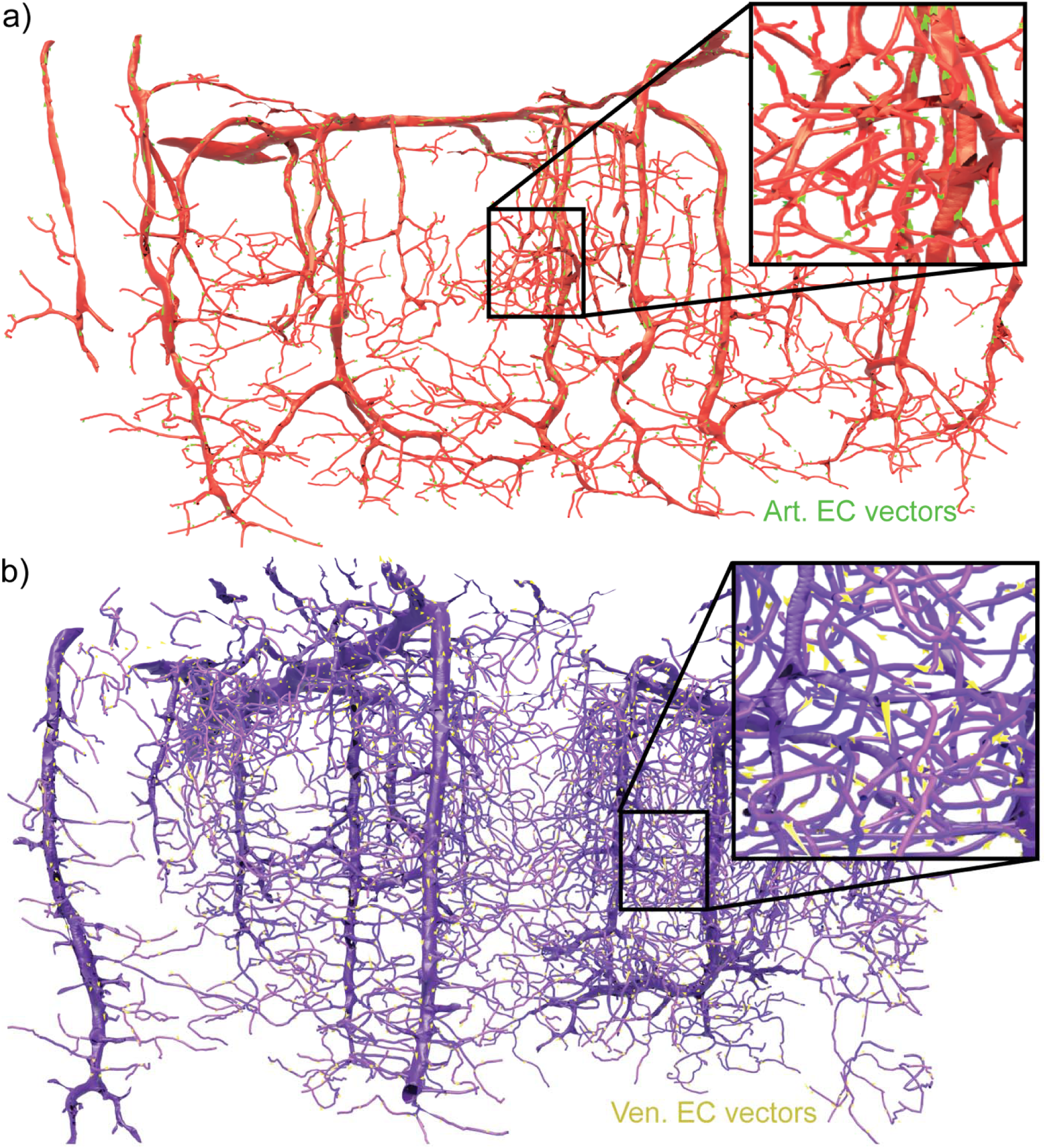
Brain microvascular endothelial cell nucleus-to-centriole distance map (a-b) Three-dimensional representations of MICrONS arterial (a, red) and venous-end (b, purple) microvasculature with EC nucleus-to-centriole distances represented as green and yellow vectors, respectively.

**Supplemental Figure 6:**
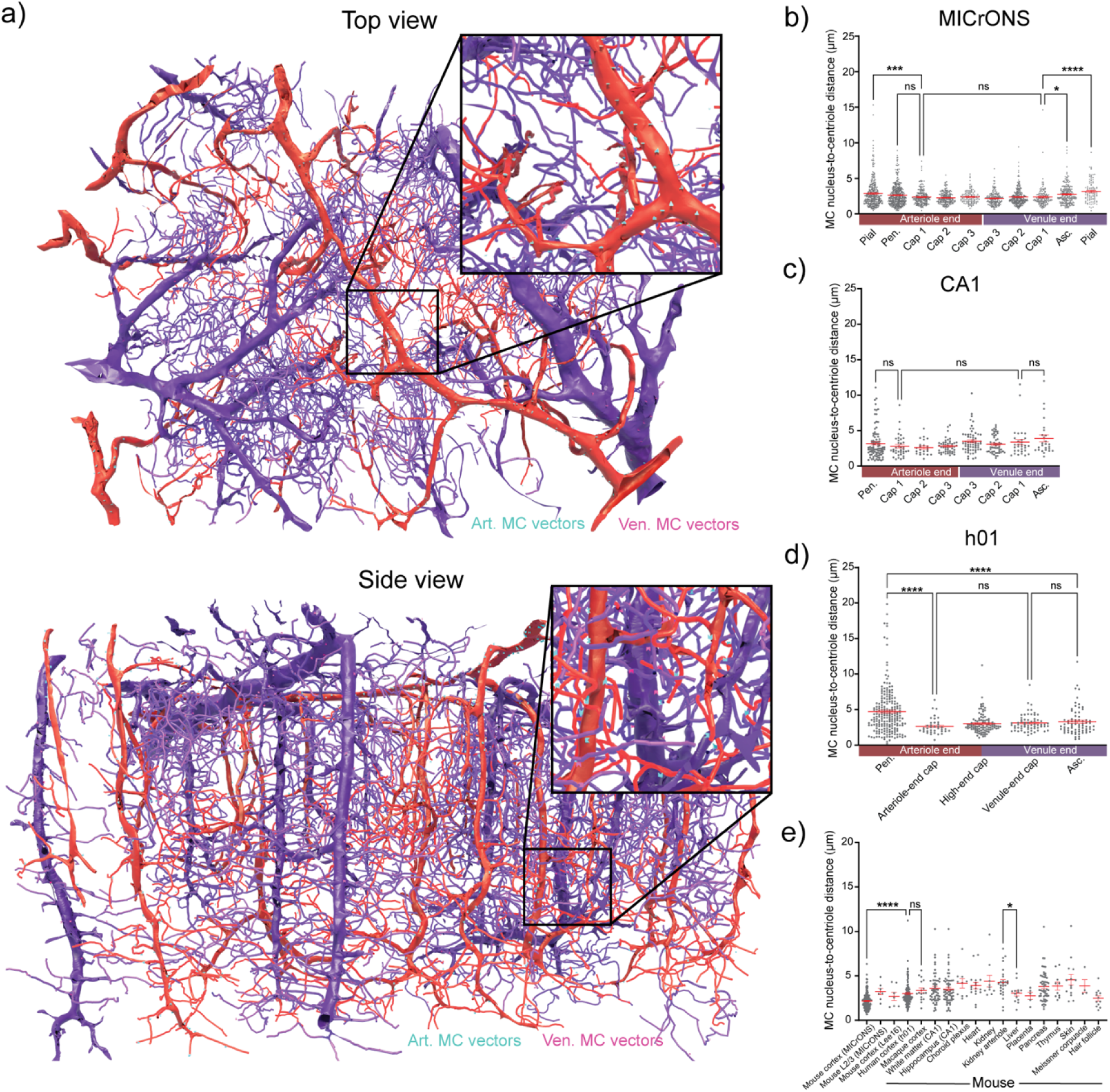
Mural cell nucleus-to-centriole distances. **(a)** Three-dimensional representations of MICrONS arterial (red) and venous-end (purple) microvasculature with MC nucleus-to-centriole distances represented as light blue and pink vectors, respectively. *Top:* top view with an enlargement as an inset. *Bottom*: side view of the MICrONS dataset. **(b-d)** MC nucleus-to-centriole lengths of arterial and venous-end microvasculature of **(b)** MICrONS, **(c)** CA1, **(d)** h01 datasets and **(e)** other brain and peripheral tissues.

## Discussion

Extensive analysis of large-volume EM datasets of the brain vasculature enabled us to systematically assess the organization of vascular cells and the distribution of their cilia and centrosomes. This strategy with nanometer resolution over continuous vascular segments, enabled precise cell identification in their native 3D milieu, which is difficult to achieve through conventional EM or light microscopy. Our findings using this approach are summarized in **Figure 7 and Sup. Video 5**. We found that a substantial number of vascular cells are ciliated, with some interspecies differences, and the prevalence of primary cilia depends on the cell subtype and vascular location. While human ECs exhibited primary cilia, we did not find a single primary cilium among the thousands of ECs we evaluated in the mouse cortex or CA1 region. Additionally, we provide the first ultrastructural evidence for the presence of primary cilia on brain MCs, with a gradient of distribution across vasculature - showing the highest frequency and greatest length in the venules and venous-end capillaries. Notably, cilia position and contact with extracellular environment differed between human and mouse datasets. Similarly, ECs density distribution between arterioles and venules varied between human and mouse datasets. Despite these interspecies differences, both human and mouse ECs shared a common feature of polarity, with their centrioles-oriented opposite to the direction of CBF. Together, these findings reveal several novel ultrastructural features of the brain microvasculature and generates new hypotheses, including a potential role of vascular primary cilia in sensing and transmitting signals between the neuropil and vasculature, as well as the involvement of venous vasculature in regulation of CBF via ciliary pathways.

**Figure 7.**
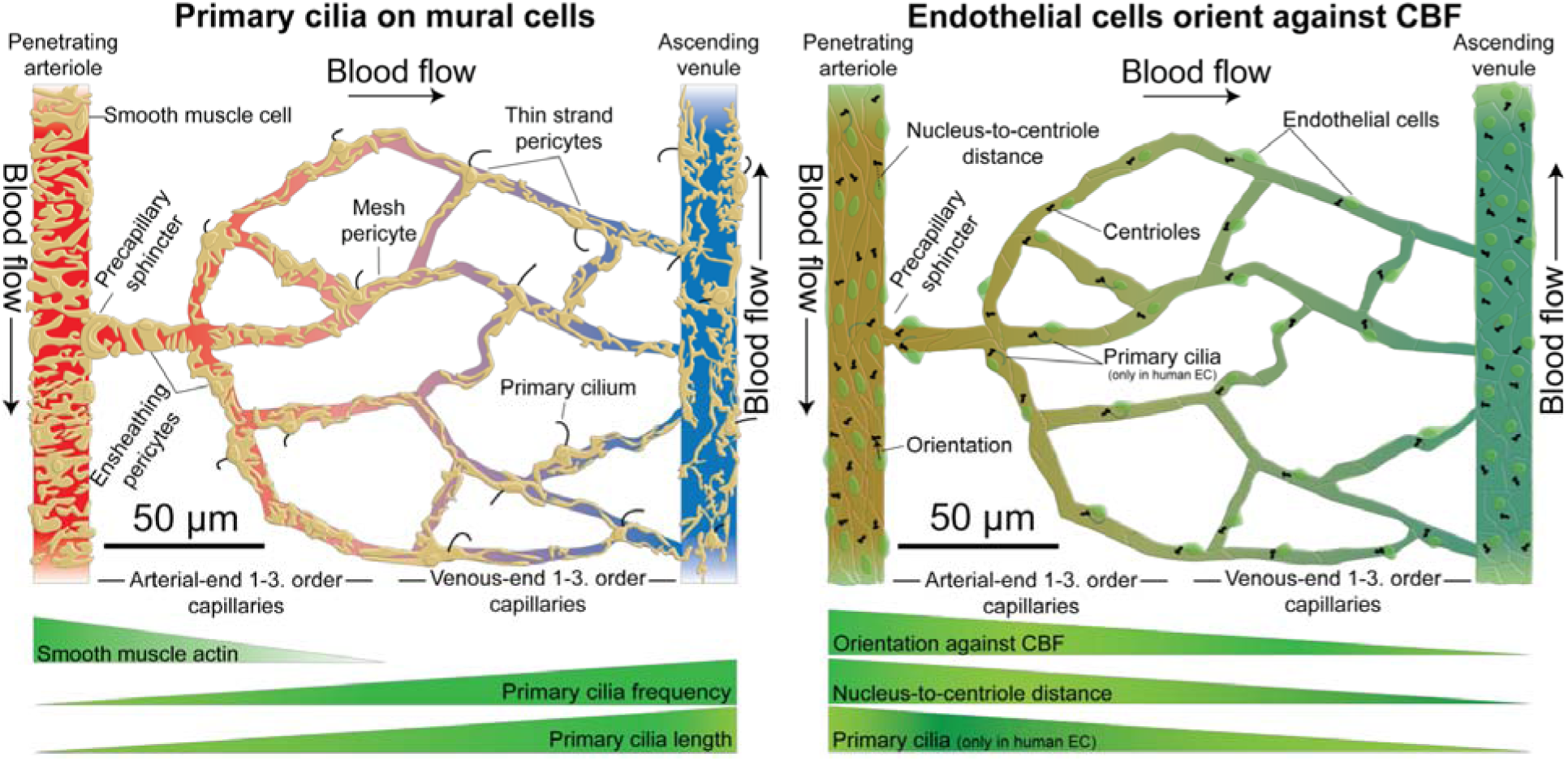
– Illustrations of the microvascular cells, centrioles and primary cilia. Graphical abstract showing gradient distribution of primary cilia in the mural cells (left) and centrioles orientation in the endothelial cells (right) in the brain microvasculature. *Left*: Primary cilium frequency and lengths are reversely proportional to smooth-muscle actin expression. *Right*: EC centrioles orientation against CBF, their nucleus-to-centriole distance, and the presence of primary cilia (in human) decreases from the arterial to venous end of the microvasculature.

In this study, we gained an unprecedented view of the cerebrovascular ultrastructure within its network context and analyzed thousands of vascular cells across species and tissues, ensuring statistical robustness of our findings. However, since only a few datasets are available for each species, some of the observed differences could reflect regional or age-related variability rather than true interspecies differences. Future large-volume EM datasets from multiple regions and developmental stages will be essential to address this. The perfusion-fixed mouse and macaque tissues were of markedly higher quality than the post-fixed human sample, which was derived from a patient and displayed astrocytic swelling, thickening of basal lamina and vessel lumen compression. Nonetheless, key ultrastructural features such as centrosome and primary cilia base positioning are unlikely to be substantially affected by these limitations.

### Primary cilia in vascular cells: Sensory antennae for neurovascular coupling

The primary cilium acts as a sensory “antenna” of the cell, capable of detecting signals from the local environment and transducing them intracellularly ^69^. The presence of primary cilia exclusively on human ECs is an unexpected finding. Whether these primary cilia sense the shear stress, oxygen levels, and blood flow, or transmit blood-borne signals to ECs remains to be evaluated. If mouse ECs truly lack primary cilia, this raises important questions about the suitability of rodent models for studying EC ciliary functions. In our study, the mouse datasets were derived from the visual cortex and CA1 region; thus, the possibility that EC primary cilia presence in other brain regions – such as the temporal cortex – cannot be excluded until the entire brain is systematically examined.

Our in-depth analysis of primary cilia in brain MCs revealed notable differences between mouse and human MC cilia. In mice, primary cilia were often ensheathed by astrocytic endfeet and occasionally penetrated between the endfeet to come in direct contact with the neuropil. In contrast, human MCs rarely formed surface cilia and their cilia typically extended along the endothelial-facing side of the MC. Interestingly, like human MCs, EC-facing mouse MC cilia were also less likely to grow directly from the cell surface compared to cilia facing the parenchyma. Overall, cilia origins in MCs were far less uniform than those reported in other cell types like neurons ^41^. Moreover, the high prevalence of primary cilia forming a partial pocket suggests that surface and pocket ciliogenesis may represent a continuum, rather than distinct, discrete processes.

Despite these interspecies differences, a striking feature is that primary cilia are preferentially located on venules and capillaries surrounding the venules in all datasets. Our understanding of the roles of brain venous vasculature remains limited. Recent studies ^10,11^ provide compelling evidence that venous vasculature may participate in regulation of CBF. However, further studies are needed to clarify the precise functional role of the venous vasculature in controlling CBF. Given the high prevalence of primary cilia on venous MCs and the well-recognized role of primary cilia as sensory organelles in other cell types, including neurons and astrocytes ^41,42,70^, this new data leads us to hypothesize that primary cilia-dependent signaling pathways in the venules contributing to NVC. Venous vasculature exhibits comparatively higher permeability than arterioles and serves as potential routes for paracellular transport and clearance of the metabolic by-products ^43,71^. Since primary cilia are hubs for ion channels, transporters, and receptors ^69^, they could sense local changes in the environment – including elevated concentrations of metabolites such as K^+^, amyloid beta, and glutamate near astrocytic endfeet and in the parenchyma – thereby potentially contributing to the regulation of BBB permeability and clearance mechanisms.

The high prevalence of primary cilia in capillary MCs (pericytes) and their proximity to astrocytes and neuronal synapses supports a potential role in NVC at the capillary level. We and others have identified multiple NVC mechanisms initiated by neuroglial mediators – such as K^+^, Gq-protein coupled receptor agonists, and adenosine – that engage capillary EC and MC molecular machinery to regulate CBF ^10,52,53,72–74^. How these signals are sensed and transmitted remains an open question. Whether primary cilia act as the sensory “antennae” receiving signals from the neuropil and transmitting it to the vasculature will need to be addressed in future animal studies. Notably, the astrocytic endfeet ensheathment and penetration to the neuropil observed in mouse MC cilia were absent in the human sample, raising doubts that cilia serve as the sole mechanism for signal detection. In the human (h01) dataset, all MC cilia were confined within the basal lamina. Extensive global structural changes have been observed in capillaries minutes after cardiac arrest (see Sup. Video 7 from ^75^). Thus, we cannot exclude the possibility that primary cilia in the human (h01) sample shifted from their native context during tissue excision and drop fixation.

The absence of primary cilia in arteriolar MCs and —many ECs — introduces another open question about the functional significance of vascular cilia. The gradient distribution pattern may reflect a specialization related to flow, pressure, or permeability differences across the vasculature. The predominance of unciliated arteriolar MCs versus the higher prevalence of cilia on venous-end MCs correlates with actin-myosin levels and, by extension, vascular reactivity - a relationship that warrants further investigation. Interestingly, primary cilia were also present on vascular MCs in multiple peripheral mouse tissues, including heart, liver, and kidney. Whether a similar arteriovenous gradient of ciliation exists in these peripheral vascular beds provides an intriguing direction for future studies. Although primary cilia are often described as nearly universal, non-ciliated cells exist throughout the body, and lack of primary cilia may be both necessary and functionally important ^76^. Expanding these analyses to human, primate, and rodent tissues from both healthy and diseased states will be essential for advancing our understanding of the role of primary cilia in vascular function. Given that many neurological disease and age-related conditions are linked to ciliopathies, our identification of ciliated vascular cells provides an important foundation for future studies exploring vascular ciliopathies as potential contributors to VCID and other neurological diseases.

### Vascular cell density represents structural differences across the brain vasculature

Our annotations of ECs and MCs across the datasets provide an unprecedented high-resolution map that we used to quantify vascular cell densities throughout the brain microvasculature. To our knowledge, this is the first extensive quantification and comparison of EC and MC density between the arteriolar and venular ends. Previous studies have reported total vessel length and vascular nuclei count in brain samples from mouse ^60^, monkey ^61^, and human ^62,63^. For example, Tsai et al. estimated approximately 0.88 m of vasculature per mm^3^ of the mouse cortex containing ∼43,000 vascular-cell nuclei ^60^, whereas the human cortex was reported to have roughly half that vessel length (∼0.2 – 0.5 m per mm^3^) ^62,63^. However, these studies did not assess vascular cell numbers or densities. Our study therefore fills a critical gap in the knowledge regarding vascular cell density and provides a valuable reference for future research on the structural and functional organization of the brain microvasculature.

Vascular ECs, together with tight junction proteins, constitute the fundamental structural components of the BBB ^59^. Given that venules have recently been shown to exhibit higher permeability ^71^, it was surprising to observe similar EC density between arteriolar and venular ends of the mouse cortex and CA1 region. In agreement with this observation, single cell RNA-seq data reveal comparable expression of tight junction proteins, including claudin-5, occludin, and zona occludens-1, in arteriolar and venous ECs ^77,78^. This raises an important question: if EC density and tight junction protein expression are similar, what accounts for the increased BBB permeability at the venous end? In contrast, the human cortex showed significantly lower EC density in venules compared to arterioles. However, given the structural deformities present in the human samples, further studies are required to determine whether this reduced ECs density is specific to the human brain or reflects sample-related artifacts.

Both ECs and MCs participate in multiple mechanisms of NVC ^10,52–55,72–74^. Several NVC pathways rely on long-distance signal propagation along the vasculature ^10,52,54,79^, and cellular and subcellular distribution is a key determinant of how such signals are transmitted ^79^. In this study, we not only document EC distribution across the vascular tree, but we also extend this analysis to MCs, thereby contributing to a deeper understanding of the cellular organization underlying vascular signal propagation ^1,80^. Moreover, we provide evidence that venules have sparse coverage of MCs compared to arterioles. These observed differences in MC density between arteriolar and venous ends align with their distinct physiological roles: higher MC density supports the well-established contractile function of arteriolar MCs, while reduced MC density at the venous end may represent a key structural determinant underlying the increased vascular permeability observed in venules. While further confirmation using comparable large volume EM datasets is required to validate the observed interspecies differences in vascular cell density, our findings provide a strong foundation for linking structural and functional mechanisms to cellular organization in the brain vasculature.

### ECs polarize against the CBF

Blood-flow shear stress is a key determinant of vascular network optimization. However, our understanding of how vascular cells pattern and polarize in response to flow direction has only recently begun to emerge ^81^. Centriole positioning establishes mammalian cell polarity ^66^ and is critical for endothelial barrier function by maintaining cell-cell contacts ^13^. In our analysis, the vast majority of EC centrioles were positioned upstream of the nucleus center relative to the direction of blood flow in penetrating arterioles, ascending venules, and capillaries. This orientation was maintained in pial arterioles but became heterogenous in larger pial venules, suggesting that flow rate and shear stress are major determinants of EC polarity ^82,83^. Consistent with our findings, EC polarity against the blood flow direction has been reported in the mouse retina using Golgi apparatus positioning as a polarity marker ^82^. In contrast, in peripheral vessels such as the rabbit inferior vena cava, and in the pig and rat aorta, centrioles were located on the heart side of the nucleus, regardless of flow direction ^84,85^. These contrasting results suggest that centriole positioning — and thus polarity — may be flow-dependent in the cerebral circulation, but not in the peripheral vasculature.

It remains unclear why brain EC centrioles consistently orient opposite to blood flow, whether this pattern is developmentally programmed and maintained in the mature cells. Although centriole positioning may be influenced by their role in ciliogenesis, our analysis revealed that centrioles oriented against flow in both mouse and human ECs, whereas primary cilia were only present in human ECs demonstrating absence of direct relationship with ciliogenesis. Altogether, centriole orientation opposite to flow may serve as a biomarker for predicting blood-flow direction and local shear stress in the brain, a finding particularly useful for identifying flow direction in volume EM or microscopy datasets with limited vascular context.

### Unanswered questions and future directions

One prominent unanswered question is the functional role of primary cilia in the brain vasculature. The presence of MC primary cilia frequency across the brain vasculature raises the possibility that these organelles have specialized functions as sensory antennae and beyond. In the mouse brain, MC primary cilia occasionally penetrate astrocyte endfeet to reach the neuropil and make close contact with neuronal synapses, suggesting a potential role in NVC. Testing these hypotheses will requires *in-vivo* experiments using conditional knockout models targeting primary ciliary proteins such as IFT88 and Arl13b. Our examination of smaller EM volumes from peripheral tissues and other brain regions also revealed the presence of ciliated MCs, warranting further investigation into whether these tissues display a similar venular-end enrichment.

The higher prevalence of primary cilia on venular-end MCs raises the possibility that primary cilia might have a role in maintaining BBB functions. In line with this idea, recent study by Shih and colleagues demonstrated that reduced capillary-to-venous drainage in white matter contributes to age-related hypoperfusion^11^. Future studies assessing primary cilia density in white matter may reveal whether impaired drainage is associated with cilia dysfunction or broader vascular ciliopathies.

Our understanding of the role of the brain’s venous vasculature in regulating CBF remains limited. The present findings — particularly the consistent orientation of centrioles as a marker of EC polarity and the comparable EC distribution across arteriolar and venous ends — underscore the need for a more detailed examination of venous vascular cells in NVC. The observation that centrioles align opposite the direction of CBF raises intriguing questions about how EC polarity is established during development and how it is altered in neurovascular disorders. Experimental models of disturbed shear stress and blood flow, including hypertension, stroke, and other small vessel diseases, such as CADASIL, will be instrumental in defining the relationship between EC polarity and vascular dysfunction. The frequent occurrence of pial vessel anastomoses and collateral formations in stroke models ^86,87^, together with the reversed centriole orientation observed across anastomoses, could provide new mechanistic insight into EC migration and BBB alterations under pathological flow conditions.

In summary, our identification of primary cilia within the brain vasculature, their spatial orientation, and their intimate association with neural cells offer new perspectives on cellular communication in the neurovascular unit. These findings open a path toward dissecting the molecular machinery by which primary cilia and centrioles coordinate vascular sensing, polarity, and hemodynamic regulation.

## Acknowledgements

We acknowledge and thank the MICrONS Consortium, Helene Schmidt (Max Planck Institute, Germany), Lichtman laboratory (Harvard University), the Connectomics (Google team), Janelia OpenOrganelle (HHMI, Virginia, USA), and Rohan Lewis (University of Southampton) for sharing the large volume electron microscopy datasets. We thank Tyler Sloan (Quorumetrix Studio) for preparing videos for the visualization.

## Author contribution

A.M. and S.G. conceptualized the project; A.M., S.G. and C.O. wrote initial draft of the manuscript.; V.C., S.G., C.O, A.M. performed annotations and analyzed the datasets; A.M., S.G., V.C., C.O. and J.L.S. edited the manuscript.

## Funding statement

This research was supported [in part] by the Intramural Research Program of the National Institutes of Health (NIH). The contributions of the NIH author(s) were made as part of their official duties as NIH federal employees, are in compliance with agency policy requirements, and are considered Works of the United States Government. However, the findings and conclusions presented in this paper are those of the author(s) and do not necessarily reflect the views of the NIH or the U.S. Department of Health and Human Services. This research was supported by the Intramural Research Program of the National Institute of Neurological Disorders and Stroke, NIH, Bethesda, MD to A.M. (NS009456-01) and Danish Research Council to S.G. (2034-00304B). C.M.O. and J.L.-S. were supported by the Howard Hughes Medical Institute.

## Conflict of interest disclosure

The authors declare no competing interests.

## Supplementary video legends

**Sup. Video 1:** Serial images from the volume EM segmented to show a capillary primary cilium ensheathed by astrocyte endfeet

**Sup. Video 2:** Serial images from the volume EM segmented to show a capillary primary cilium penetrating into the neuropil

**Sup. Video 3:** Segmented brain microvasculature from MICrONS dataset shows that primary cilia are selectively located around the venules and capillaries

**Sup. Video 4:** Segmented brain microvasculature from MICrONS dataset including nucleus to centriole vectors in endothelial cells.

**Sup. Video 5:** Video summarizing the findings showing cilia and centriole organization across the microvasculature in the MICrONS dataset.

## Notes

### Competing Interest Statement

The authors have declared no competing interest.

